# Immunogenic SARS-CoV2 Epitopes Defined by Mass Spectrometry

**DOI:** 10.1101/2021.07.20.453160

**Authors:** Ke Pan, Yulun Chiu, Eric Huang, Michelle Chen, Junmei Wang, Ivy Lai, Shailbala Singh, Rebecca Shaw, Michael MacCoss, Cassian Yee

## Abstract

SARS-CoV-2 infections elicit both humoral and cellular immune responses. For the prevention and treatment of COVID19, the disease caused by SARS-CoV-2, T cell responses are important in mediating recovery and immune-protection. The identification of immunogenic epitopes that can elicit a meaningful T cell response can be elusive. Traditionally, this has been achieved using sophisticated *in silico* methods to predict putative epitopes; however, our previous studies find that ‘immunodominant’ SARS-CoV-2 peptides defined by such *in silico* methods often fail to elicit T cell responses recognizing SARS-CoV-2. We postulated that immunogenic epitopes for SARS-CoV-2 are best defined by directly analyzing peptides eluted from the peptide-MHC complex and then validating immunogenicity empirically by determining if such peptides can elicit T cells recognizing SARS-CoV-2 antigen-expressing cells. Using a tandem mass spectrometry approach, we identified epitopes of SARS-CoV-2 derived not only from structural but also non-structural genes in regions highly conserved among SARS-CoV-2 strains including recently recognized variants. We report here, for the first time, several novel SARS-CoV-2 epitopes from membrane glycol-protein (MGP) and non-structure protein-13 (NSP13) defined by mass-spectrometric analysis of MHC-eluted peptides, provide empiric evidence for their immunogenicity to induce T cell response.

**Significance Statement:** Current state of the art uses putative epitope peptides based on *in silico* prediction algorithms to evaluate the T cell response among COVID-19 patients. However, none of these peptides have been tested for immunogenicity, i.e. the ability to elicit a T cell response capable of recognizing endogenously presented peptide. In this study, we used MHC immune-precipitation, acid elution and tandem mass spectrometry to define the SARS-CoV-2 immunopeptidome for membrane glycol-protein and the non-structural protein. Furthermore, taking advantage of a highly robust endogenous T cell (ETC) workflow, we verify the immunogenicity of these MS-defined peptides by in vitro generation of MGP and NSP13 peptide-specific T cells and confirm T cell recognition of MGP or NSP13 endogenously expressing cell lines.

## Introduction

Severe acute respiratory syndrome coronavirus 2 (SARS-CoV-2), the highly transmissible respiratory virus responsible for the COVID-19 pandemic outbreak, continues to render significant, lasting impact on global public health and has created an urgent need to develop accurate immunodiagnostics, and effective treatment strategies (1, 2). Rapid dissemination of the SARS-CoV-2 genomic sequence first revealed by Dr. Zhang Yongzhen led to large scale efforts around the world to develop a protective vaccine that could elicit humoral (antibody) and cellular (T cell) responses (3). It follows that the identification of immunogenic epitopes of SARS-CoV-2 recognized by the human immune system would be critical for rational vaccine development.

Using *in silico* prediction algorithms, several investigators have amassed extensive panels of Class I and Class II restricted epitopes to probe SARS-CoV-2-specific T cell responses, in some cases, combining these with overlapping ‘megapools’ spanning regions conserved regions of the genome (4, 5). These peptides have been used to track responses in infected and convalescent individuals (6, 7), design multi-epitope vaccines and used directly or indirectly to measure the breadth and severity of COVID19 disease (7–13). While these studies have uncovered insights into the T cell immunobiology of COVID19, the accuracy of T cell responses using in silico predicted responses and overlapping long peptide (OLP) pools is diminished by a failure to consider whether such epitopes are immunogenic. An immunogenic epitope in this sense is defined as a peptide that is known to be presented by self-MHC, and is capable of eliciting T cells of sufficient affinity that such T cells can recognize target cells endogenously expressing antigen and presenting the antigen-derived peptide in the context of an MHC complex with sufficient surface density as to sensitize the target cell to peptide-specific T cell-mediated recognition. In essence, an immunogenic epitope of SARS-CoV-2 requires both direct sequencing of peptides presented by MHC as well as empiric validation of T cell immunogenicity.

To our knowledge, this study is the first use of tandem MS to identify T cell epitopes of SARS-CoV-2 conserved protein, MGP and NSP13, following peptide elution from the MHC complexes of SARS-CoV-2-expressing cells, and the first study to empirically validate immunogenicity by in vitro generation of SARS-CoV-2-specific CTL. Enabling technology developed by our group for the isolation of rare tumor-reactive T cells from very low precursor frequency populations in the peripheral blood was applied (14). We present data on the identification of five immunogenic epitopes of a highly conserved region of MGP and the NSP13 region of the SARS-CoV-2 genome and demonstrate that such MGP65- and NSP13-specific CTL recognize and kill SARS-CoV-2 antigen expressing target cells.

## Results

### Profiling of MHC Class-I restricted epitope of SARS-CoV-2

The antigen discovery platform for SARS-CoV-2 is comprised of four steps: (1) peptide elution and identification with mass spectrometry (MS) for SARS-CoV-2 targets; (2) endogenous T cell (ETC) generation workflow to elicit peptide-specific CTL; (3) empiric validation of antigen specific CTL against SARS-CoV2 targets; and (4) SARS-CoV-2 specific T cell receptor (TCR) engineered T cell (TCR-T) development (Fig. S1). In order to elute and sequence the MHC bound peptide derived from SARS-CoV-2, the SARS-CoV-2 genes were overexpressed in targets cells with different HLA allele expression. Lentiviral expression vectors spanning highly conserved regions of SARS-CoV-2 regions: membrane glycol-protein (MGP) or Non-structure protein helicase (NSP13)(10, 23), were constructed and used to infect target cell line A375 (HLA-A0101/0201), Mel624 (HLA-A0201), RPMI-7951 (HLA-A0101/0201), Hs-578T (HLA-A0301/A2402) and M14 (HLA-A1101/2402). Following puromycin selection and expansion of MGP or NSP13 stably -expressing cell lines, purity of over 90% was achieved (data not showed). The MGP or NSP13-expressing cell lines were expanded to 300 – 500 million, harvested, lysed with NP40 detergent lysis buffer and subjected to total HLA class I immunoprecipitation (anti-HLA-A, B, C) and acid elution, followed by tandem mass spectrometry (MS) to analyze the HLA-bound peptides.

We initially analyzed the eluted HLA bound peptides derived from the SARS-CoV-2 targets established above by using data dependent analysis liquid chromatography tandem mass spectrometry (DDA MS/MS). The eluted spectra were searched using the Mascot search engine node (version 2.6) within Proteome Discoverer (version 2.3) processing workflow with the Swiss-Prot human proteome database (version 2020_05) followed by virus proteome database (version 2020_05). To reduce false positive hits from human proteome, the “Spectrum Confidence Filter” node within Proteome Discoverer processing workflow filtered out all spectra with highly confident peptide-spectrum matches annotated from the human proteome. The remaining spectra were further searched against the virus proteome (Fig. 1A). In total, 12,770 MS/MS were acquired, 9731 Peptide-Spectrum-Match (PSM) were annotated yielding 357 peptides with Mascot Ions Score ≥ 25. Among the 357 peptides was annotated a peptide derived from NSP13 (NSP13-400, VYIGDPAQL) with Mascot Ions Score=27 (Fig. 1B). This peptide was eluted from M14-NSP13 cells (HLA-A1101/A2402). From the HLA binding prediction using IEDB tool, the NSP13-400 peptide scored high predicted binding affinity to HLA-A2402 allele (Table 1), suggesting that NSP13-400 peptide is likely to be presented by HLA-A2402.

**Fig. 1.**
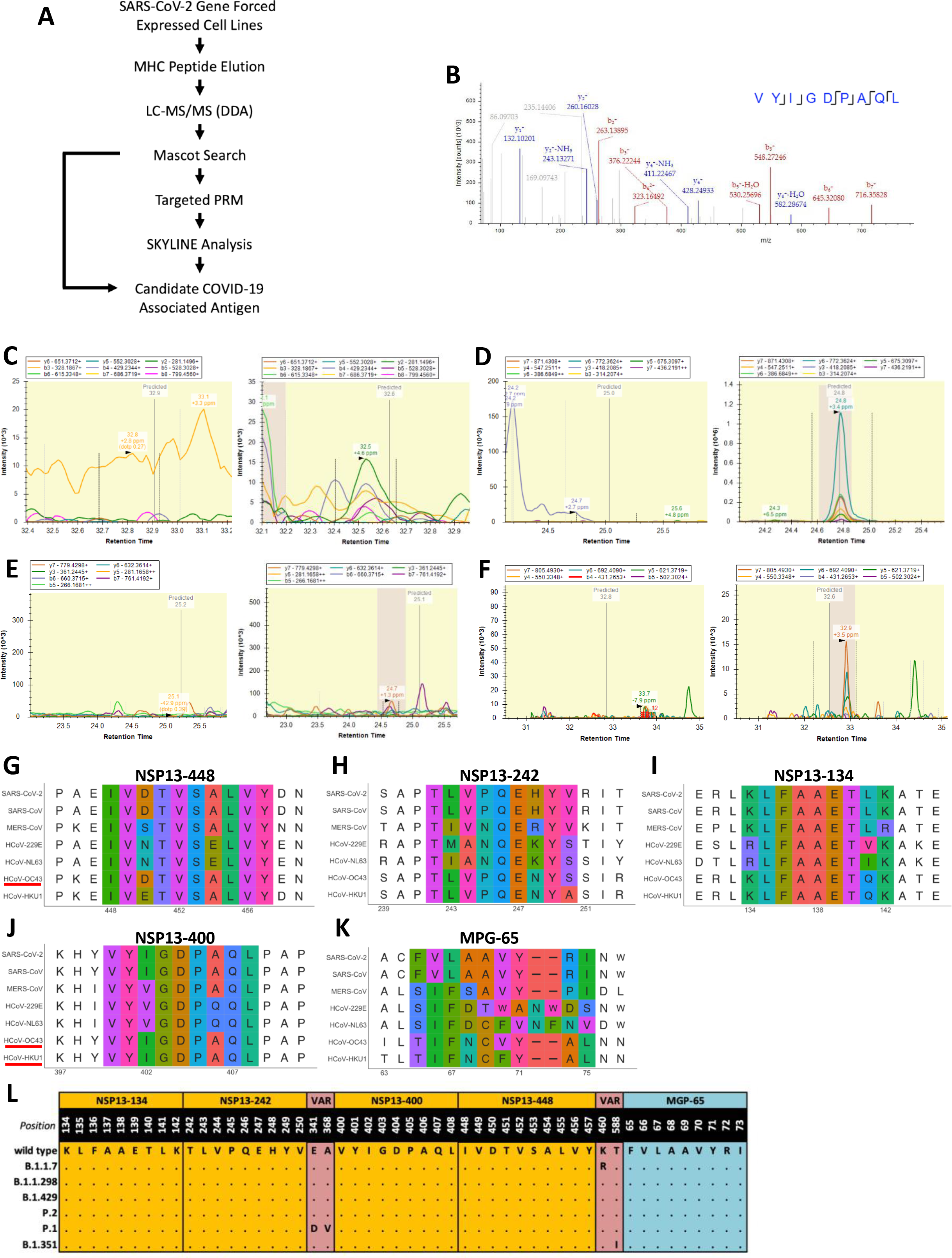
SARS-CoV-2 derived HLA Class-I peptide identification with Mass Spectrometry. (A) Schematic representation of HLA-I peptide identification. MS/MS spectra of immunopeptidome and proteome analysis were searched against with Swiss-Prot human and virus protein database and filtered at 1% FDR. (B) MS/MS annotation for NSP13-400 peptide (VYIGDPAQL). (C-F) PRM analysis for NSP13-448 peptide (IVDTVSALVY), NSP13-242 peptide (TLVPQEHYV) NSP13-134 peptide (KLFAAETLK) and MGP-65 peptide (FVLAAVYRI). MS1 XIC areas and MS/MS for each targeted peptide were plotted using Skyline software. (G-K) The multiple sequence alignment of the four candidate peptide sequences to all coronavirus known to infect humans. (L) Four mutations were reported from NSP13 protein: E341D, A368V (P.1 lineage), K460R (B.1.1.7 lineage) and T588I (B.1.351 lineage). The following abbreviations are used: VAR: variants, a dot (.) indicates the same amino acid in that position.

**Table 1.**
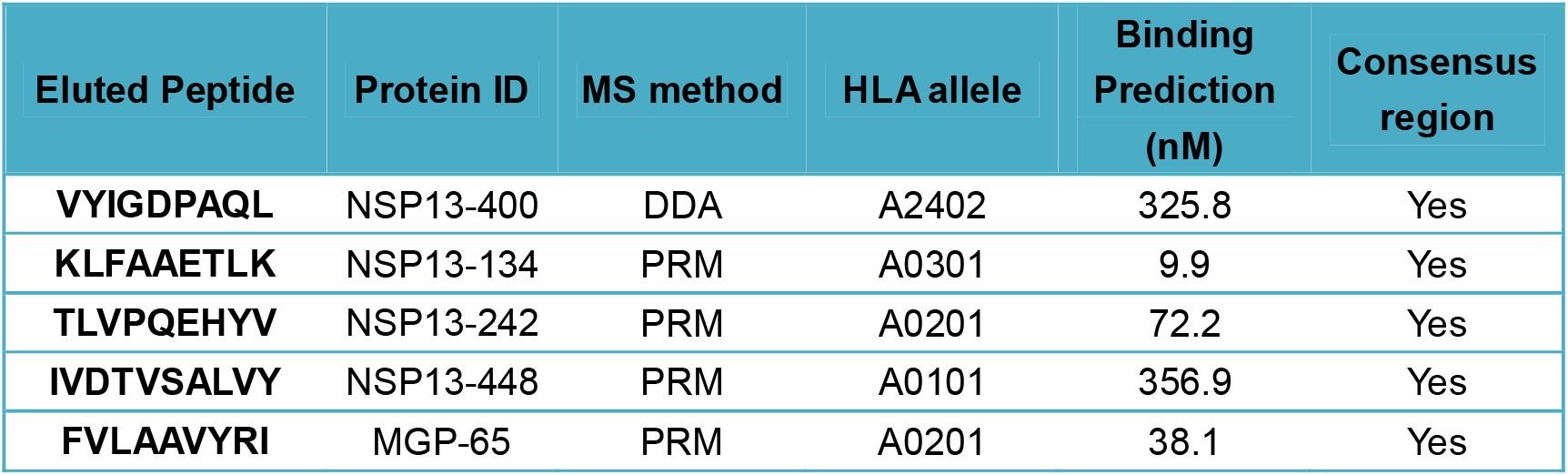
Summary of identified SARS-CoV-2 epitopes.

To enable more comprehensive profiling of potential HLA Class-I restricted peptides from SARS-CoV-2, we further analyzed eluted peptides by parallel reaction monitoring mass spectrometry (PRM-MS) to focus predicted high probability HLA binding peptides derived from SARS-CoV-2 but not successfully detected by DDA approach. Prior to the PRM-MS, ten predicted high potential HLA-A0101, HLA-0201 or HLA-A0301 binding peptides from MGP or NSP13 were selected, respectively (Table S1). For the eluted peptide from target cell lines, precursor ion inclusion lists of 10 potential peptides were generated using Skyline, and we targeted and monitored these 10 peptides using nanoflow LC-PRM-MS with high mass accuracy and resolution. Pierce™ Peptide Retention Time Calibration Mixture peptides were used to monitor retention time drifts and adjust the scheduled PRM method. We first generated spectral library using synthetic peptides, and we used both synthetic peptides and Pierce™ Peptide Retention Time Calibration Mixture peptides to define iRT, a normalized dimensionless peptide-specific value, to accurately predict retention time of each targeted peptide. We detected IVDTVSALVY (NSP13-448) with Dotp=0.58 and average product ion ppm error at +4.6 ppm at predicted time 32.6 min (Fig. 1*C* Right) in target cell line RPMI-7951-NSP13 and no peak was detected in the negative control cell line RPMI-7951-GFP at predicted time 32.9 min (Fig. 1*C* Left). TLVPQEHYV (NSP13-242) with Dotp = 0.76 and average product ion ppm error at +3.4 ppm at predicted time 24.8 min (Fig. 1*D* Right) was detected at target cell line A375-NSP13 and no peak was detected in the negative control cell line A375-GFP at predicted time 25.9 min (Fig. 1*D* Left). KLFAAETLK (NSP13-134) with Dotp=0.67 and average product ion ppm error at +1 ppm at predicted time 25.1 min was detected in target cell line Hs-578T-NSP13 (Fig. 1*E* Right) and no peak was detected in the negative control cell line Hs-578T-GFP at predicted time 25.2 min (Fig. 1*E* Left). Same rules applied. From A375-MGP, we detected FVLAAVYRI (MGP-65) with Dotp=0.91 and average product ion ppm error at +2.5 ppm at the predicted time 32.7 min (Fig. 1*F* Right) and no peak was detected in the negative control cell line A375-GFP at predicted time 32.8 min (Fig. 1*F* Left). These XIC MS2 analyses reported sufficient well-defined peaks in the positive control and no peaks showing in the negative control (matrix blank), suggesting that these targeted peptides exist in eluted peptide samples.

To evaluate if these five candidates SARS-CoV-2 HLA Class-I restricted peptides identified with DDA or PRM-MS are homologous to other coronaviruses including SARS-CoV, Middle East respiratory syndrome coronavirus (MERS-CoV), as well as other four coronavirus 229E, NL63, OC43 and HKU1, multiple sequence alignment (MSA) analysis was performed. NSP13-242 (Fig. 1*H*), NSP13-134 (Fig. 1*I*), and MGP-65 (Fig. 1*K*) show a high degree of homology to the sequence of SARS-CoV. NSP13-400 (Fig. 1*J*) shows a high degree of homology to the sequence of SARS-CoV, HCoV-OC43 and HCoV-HKU1 (underlined in red). NSP13-448 (Fig. 1*G*) shows a high degree of homology to the sequence of SARS-CoV, HCoV-OC43 (underlined in red). In order to evaluate if these five candidates are homologous to non-coronaviruses species, the peptides were analyzed by using BLAST searches to identify the all potential source proteins. The Top 250 hits for each target sequence reported up to 88.99% identity or 100% identity but coverage up to 88.99% or 100% identity to related-coronavirus, indicated these five candidate peptide are only homologous with coronavirus but no other species (Table S2).

### Newly-defined epitopes are found in highly conserved regions of SARS-CoV-2 and SARS-CoV2 variants

Olvera *et. al* recently described the development of a COVID19 vaccine using the overlapping of SARS-CoV-2 consensus sequence (24). This paper utilized an entropy-based calculation on more than 1700 viral genome entries in NCBI and encompassed all described SARS-CoV-2 open reading frames (ORF), including recently described frame-shifted and length variant ORF. The Nextstrain project (http://nextstrain.org), an open-source project that provides a continually-updated view of publicly available data alongside powerful analytic and visualization tools to aid epidemiological understanding and improve outbreak response, provides a means to analyze genetic diversity across the SARS-CoV-2 genome. Using both these sources, we verified that these five peptides were located in a highly conserved region of SARS-CoV-2 genome (Fig. S2). Recently, genetic variants of SARS-CoV-2 have emerged on a global scale, e.g., mutation 23403A>G-(D614G) located on the spike protein, believed to render SARS-CoV-2 more infectious (25, 26). To investigate if recently reported variants are located within these five candidates SARS-CoV-2 HLA Class-I restricted epitopes, we focused on variants of concern first described in the United Kingdom (B.1.1.7), Denmark (B.1.1.298), United States (B.1.429), Brazil and Japan (P.2 and P.1), and South Africa (B.1.351). For NSP-13 region, there are two mutations, E341Y and A368V, present in P.1 variants, one mutation K460R present in B.1.1.7 variant, one mutation T588I present in B.1.351 variant, none of which overlap with these four NSP-13 HLA Class-I restricted peptides. No variant strain has been reported from the membrane glycol-protein (MGP) region (Fig. 1*L*).

### MGP-65 peptide specific cytotoxic T cells generated from the peripheral blood recognize SARS-CoV-2-MGP-expressing target cells

Following leukapheresis, HLA-A0201 healthy donor PBMC were stimulated with MGP-65 peptide (FVLAAVYRI)-pulsed autologous DCs. After two rounds of stimulation, MGP-65-A2 tetramer-positive staining populations were detected (Fig. 2*A*). About 33 wells from one 48 well plate showed clear MGP-65 peptide tetramer positive CD8+ T cell population (Fig. S3), indicating that MGP-65 peptide specific T cells are easily expanded with cognate peptide stimulation, even in the PBMC of healthy donors without a history of SARS-CoV-2 infection. Expanded MGP-65 CTLs were tested functionally using standard ^51^Cr release assays (CRA). T2 cells (HLA-A0201), pulsed with titrated amounts of MGP-65 peptide, elicited CTL recognition and killing at peptide concentrations as low as 10 pM (Fig. 2*B*), indicating very high recognition affinity of MGP-65 CTLs for cognate peptide.

**Fig. 2.**
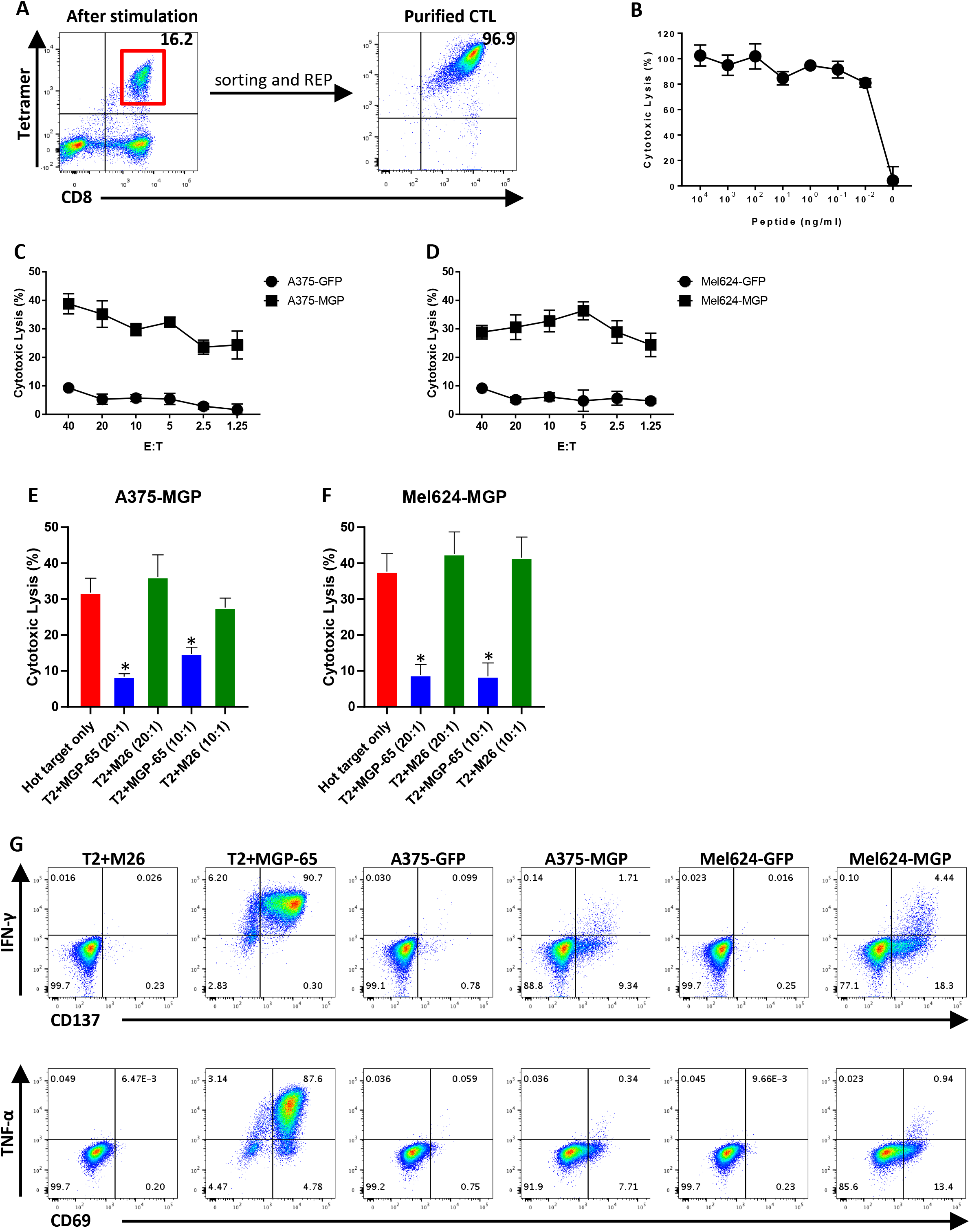
T cell generation and functional validation of membrane glycol-protein (MGP) derived HLA-A0201 peptide MGP-65. (A) Mature dendritic cells (DCs) derived from HLA-A0201 healthy donor were pulsed with MGP-65 peptide (FVLAAVYRI) and co-cultured with autologous PBMC. After two rounds of stimulation, small CD8+ and MGP-65 Tetramer+ population were observed (left side). CD8+ and MGP-65 tetramer-positive cells were then sorted and expanded using a standard rapid expansion protocol (REP). After expansion for two weeks, high purity CTLs (tetramer+ population over 90%) were generated (right side). (B) ^51^Cr labeled T2 cells pulsed with various concentrations of MGP-65 peptide were co-cultured with MGP-65 specific CTLs at 20:1 of effector to target (E:T) ratio. The lysis ability of MGP-65 specific CTLs was detected with standard ^51^Cr release assay (CRA). The data were show as average of triplicate. (C, D) ^51^Cr labeled MGP or GFP force expressing HLA-A0201 cell lines A375 (A375-MGP, A375-GFP) and Mel624 (Mel624-MGP, Mel624-GFP) were co-cultured with MGP-65 specific CTLs at various E:T ratio (from 40:1 to 1.25:1). The lysis ability of MGP-65 specific CTLs to different targets were detected with standard CRA. The data were show as average of triplicate. (E, F) Cold target inhibition assay. ^51^Cr labeled A375-MGP and Mel624-MGP were as hot targets. Non-radiolabeled T2 cells pulsed with MGP-65 peptide or M26 irrelevant peptide were as cold targets. Cold target: Hot target ratio was 10:1 or 20:1. MGP-65 specific CTLs were co-cultured with hot targets alone or hot targets together with cold targets at 20:1 E: hot T ratio. The lysis ability of MGP-65 specific CTLs were detected with standard CRA. The data were show as average of triplicate. (G) Intracellular cytokine staining (ICS) assay. MGP-65 specific CTLs were co-culture with T2 pulsed with MGP-65 peptide or M26 irrelevant peptide, as well as A375-MGP, A375-GFP, Mel624-MGP, Mel624-GFP at 10:1 E:T ratio in the presence of Brefeldin A (BFA) for overnight. After incubation, the levels of IFN-γ and TNF-α, as well as TCR pathway down-stream activated marker CD137 and CD69 were detected using flow cytometry.

To verify that MGP-65 specific CTLs recognized the endogenously presented cognate peptide, HLA-A0201+ target cells were engineered to express the SARS-CoV-2 MGP gene (A375-MGP, Mel624-MGP). MGP-65 specific CTL were able to lyse A375-MGP and Mel624-MGP cell lines, but not A375-GFP and Mel624-GFP control cell lines (Fig. 2 *C* and *D*). To further confirm that target recognition of MGP-65 specific CTL was through engagement of endogenously presented cognate peptide, cold target inhibition assay was performed. ^51^Cr pulsed MGP-expressing cells were radiolabeled with ^51^Cr while MGP-65 or M26 irrelevant peptide pulsed T2 cells were left unlabeled and used as cold targets or control cold targets, respectively. When adding cold targets at both 20:1 and 10:1 cold to hot target (C:H) ratio, the cytotoxicity of MGP-65 specific CTL for radiolabeled MGP-65 peptide targets was significantly inhibited (Fig. 2 *E* and *F*). However, there was no inhibition if control cold targets were added, indicating MGP-65 CTLs were able to then lyse MGP-65 targets via recognition of endogenously presented cognate peptide. This data provided further evidence that MGP-65 peptide is the natural endogenously presented MHC peptide.

To further evaluate function of the MGP-65 specific CTLs, intra-cellular staining (ICS) assay was performed to detect IFN-γ and TNF-α production. Co-culture of MGP-65 specific CTLs with MGP-65 peptide pulsed or MGP engineered target cells demonstrated specific recognition by IFN-γ and TNF-α produced compared with control targets pulsed with irrelevant peptide or engineered to express control GFP (Fig. 2*G*). A commensurate increase in T cell activation markers, CD137 and CD69, were also specifically and significantly elevated compared with the control group (Fig. 2*G*), when encountering relevant SARS-CoV-2 targets.

### NSP13-242-peptide specific cytotoxic T cells generated from the peripheral blood recognize SARS-CoV-2-NSP13-expressing target cells

In contrast to structural proteins such as MGP and Spike protein, non-structural proteins of SARS-CoV-2 have a lower likelihood of inducing humoral responses and neutralizing antibodies as they are not expressed on the virion surface. However, non-structural proteins of SARS-CoV-2 infected cells can be presented as MHC bound peptides and induce cellular immune responses which can be long-lasting. Here, using the same workflow, the HLA-A0201 restricted peptide, NSP13-242 (TLVPQEHYV) derived from NSP13 helicase was identified by MS/ MS. Similar to MGP-65, NSP13-242 specific T cells were readily generated using our ETC workflow (Fig. 3*A*). Surprisingly, after stimulation with NSP13-242 peptide, all 48 wells from one 48 well plates showed clear NSP13-242 peptide tetramer positive CD8+ T cell population (Fig. S4), suggesting that NSP13-242 peptide may be highly immunogenic.

**Fig. 3.**
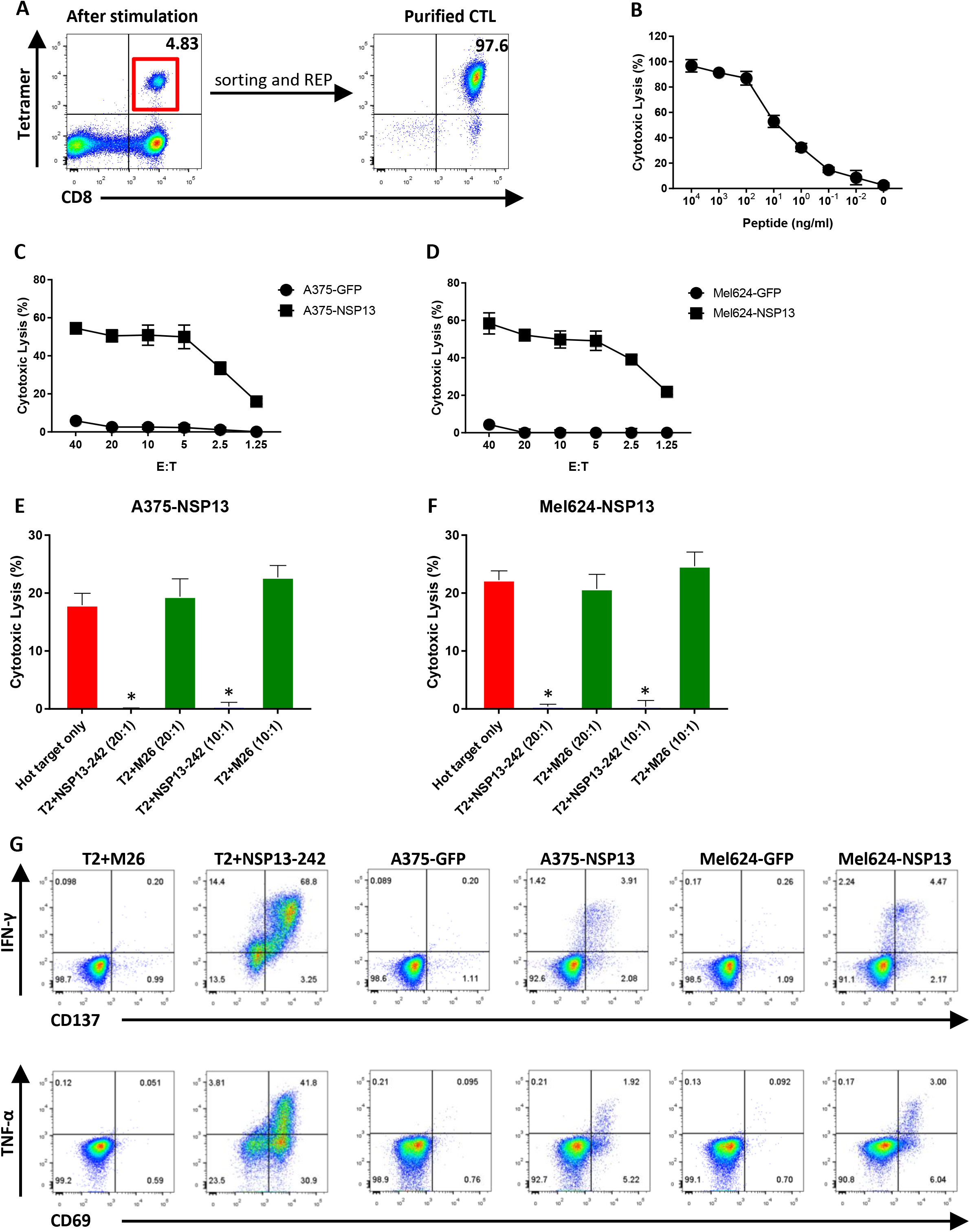
T cell generation and functional validation of non-structure protein NSP13 derived HLA-A0201 peptide NSP13-242. (A) PBMC from HLA-A0201 healthy donor were co-cultured with NSP13-242 peptide (TLVPQEHYV) pulsed autologous DCs. After two rounds of stimulation, CD8+ and NSP13-242 tetramer-positive T cells were induced (left side). CD8+ and Tetramer+ T cells were sorted and then expanded with REP for two weeks to generate high purity of NSP13-242 specific CTLs (right side). (B) ^51^Cr labeled T2 cells pulsed with various concentrations of NSP13-242 peptide were co-cultured with NSP13-242 specific CTLs at 20:1 E:T ratio. The lysis ability of NSP13-242 specific CTLs was detected with standard CRA. The data were show as average of triplicate. (C, D) ^51^Cr labeled NSP13 or GFP force expressing HLA-A0201 cell lines A375 (A375-NSP13, A375-GFP) and Mel624 (Mel624-NSP13, Mel624-GFP) were co-cultured with NSP13-242 specific CTLs at various E:T ratio (from 40:1 to 1.25:1). The lysis ability of NSP13-242 specific CTLs to different targets were detected with standard CRA. The data were show as average of triplicate. (E, F) Cold target inhibition assay. ^51^Cr labeled A375-NSP13 and Mel624-NSP13 were as hot targets. Non-radiolabeled T2 cells pulsed with NSP13-242 peptide or M26 irrelevant peptide were as cold targets. Cold target: Hot target ratio was 10:1 or 20:1. NSP13-242 specific CTLs were co-cultured with hot targets alone or hot targets together with cold targets at 20:1 E: hot T ratio. The lysis ability of NSP13-242 specific CTLs were detected with standard CRA. The data were show as average of triplicate. (G) Intracellular cytokine staining (ICS) assay. NSP13-242 specific CTLs were co-culture with T2 pulsed with NSP13-242 peptide or M26 irrelevant peptide, as well as A375-NSP13, A375-GFP, Mel624-NSP13, Mel624-GFP at 10:1 E:T ratio in the presence of BFA for overnight. After incubation, the levels of IFN-γ and TNF-α, as well as TCR pathway down-stream activated marker CD137 and CD69 were detected using flow cytometry.

Cytotoxicity assay also demonstrated that NSP13-242 specific CTLs were able to recognize cognate peptide as low as 100pM (Fig. 3*B*), indicating expression of high affinity TCR. More importantly, NSP13-242 specific CTLs were able to lyse NSP13 expressing targets A375-NSP13 and Mel624-NSP13, even at very low E:T ratio (2.5:1), but not control targets (Fig. 3 *C* and *D*), indicating that NSP13-242 specific CTLs can recognize the endogenous presented peptide of NSP13 protein. Similar to MGP-65 CTL, cold target inhibition assay also showed that when cold targets are added, the lytic capacity of NSP13-242 specific CTLs to hot targets, A375-NSP13 and Mel624-NSP13, was inhibited significantly (Fig. 3 *E* and *F*), further confirming that NSP13-242 specific CTLs lyse the targets via recognition of endogenously presented cognate peptide.

ICS assay demonstrated that NSP13-242 specific CTLs produce higher level of inflammatory cytokine IFN-γ and TNF-α and express higher levels of antigen-driven activation markers CD137 and CD69 when co-cultured with NSP13-242 peptide pulsed targets or NSP13 expressing targets, compared with the control targets (Fig. 3*G*). Thus, similar to MGP-65 specific CTLs, NSP13-242 specific CTLs will also initiate specific cellular immune response when encountering SARS-CoV-2.

### NSP13-448 peptide specific cytotoxic T cells generated from the peripheral blood recognize SARS-CoV-2-NSP13-expressing target cells

HLA-A0201 allele is expressed in about 45% of the Caucasian and Asian population (27). Specific T cell targeting of other highly prevalent HLA-A alleles of SARS-CoV-2 would be desirable given the global reach of COVID19. Using the same workflow for MGP65 and NSP13-242 peptide, we identified an HLA-A0101 restricted peptide, NSP13-448 (IVDTVSALVY) derived NSP13 protein by MS/MS; this allele covers about 26% of Caucasian and 7% of Asian population (27). Similar to NSP13-242, NSP13-448 specific T cells were readily generated using our ETC workflow (Fig 4*A*). However, after stimulation with NSP13-448 peptide, only one well from one 48 well plates showed clear NSP13-448 peptide tetramer-positive CD8+ T cell population (Fig S5), suggesting that NSP13-448 peptide may not be highly immunogenic.

**Fig. 4.**
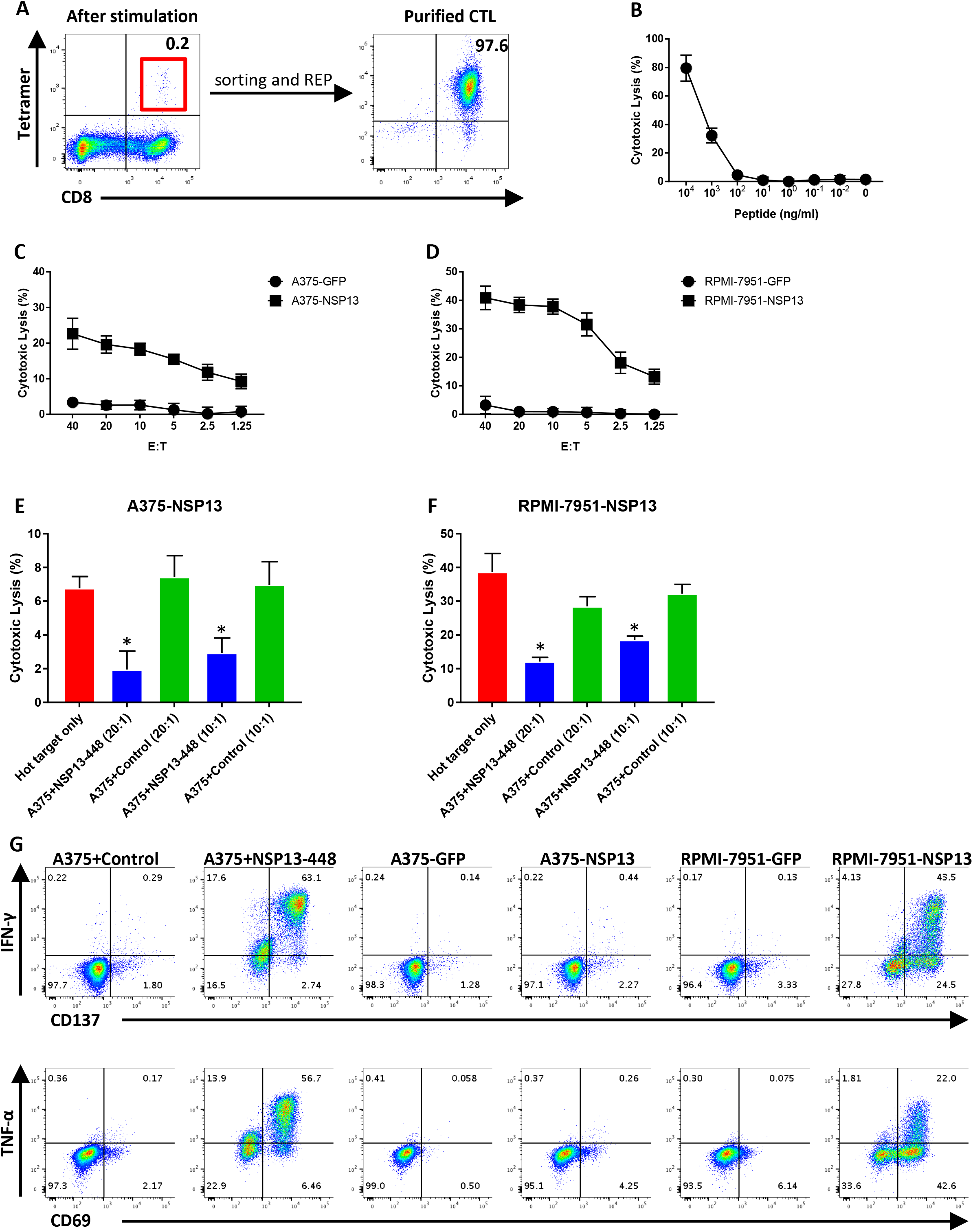
T cell generation and functional validation of non-structure protein NSP13 derived HLA-A0101 peptide NSP13-448. (A) PBMC from HLA-A0101 healthy donor were co-cultured with NSP13-448 peptide (VYIGDPAQL) pulsed autologous DCs. After two rounds of stimulation, CD8+ and NSP13-448 tetramer-positive T cells were induced (left side). CD8+ and Tetramer+ T cells were sorted and then expanded with REP for two weeks to generate high purity of NSP13-448 specific CTLs (right side). (B) ^51^Cr labeled A375 cells pulsed with various concentrations of NSP13-448 peptide were co-cultured with NSP13-448 specific CTLs at 20:1 E:T ratio. The lysis ability of NSP13-448 specific CTLs was detected with standard CRA. The data were show as average of triplicate. (C, D) ^51^Cr labeled NSP13 or GFP force expressing HLA-A0101 cell lines A375 (A375-NSP13, A375-GFP) and RPMI-7951 (RPMI-7951-NSP13, RPMI-7951-GFP) were co-cultured with NSP13-448 specific CTLs at various E:T ratio (from 40:1 to 1.25:1). The lysis ability of NSP13-448 specific CTLs to different targets were detected with standard CRA. The data were show as average of triplicate. (E, F) Cold target inhibition assay. ^51^Cr labeled A375-NSP13 and RPMI-7951-NSP13 were as hot targets. Non-radiolabeled A375 cells pulsed with NSP13-448 peptide or irrelevant HLA-A0101 peptide were as cold targets. Cold target: Hot target ratio was 10:1 or 20:1. NSP13-448 specific CTLs were co-cultured with hot targets alone or hot targets together with cold targets at 20:1 E: hot T ratio. The lysis ability of NSP13-448 specific CTLs were detected with standard CRA. The data were show as average of triplicate. (G) Intracellular cytokine staining (ICS) assay. NSP13-448 specific CTLs were co-culture with A375 pulsed with NSP13-448 peptide or irrelevant HLA-A0101 peptide, as well as A375-NSP13, A375-GFP, RPMI-7951-NSP13, RPMI-7951-GFP at 10:1 E:T ratio in the presence of BFA for overnight. After incubation, the levels of IFN-γ and TNF-α, as well as TCR pathway down-stream activated marker CD137 and CD69 were detected using flow cytometry.

Cytotoxicity assays also demonstrated that NSP13-448 specific CTLs were able to recognize cognate peptide as low as 100nM (Fig. 4*B*), indicating moderate to low TCR affinity. Interestingly, NSP13-448 specific CTLs were still able to lyse NSP13 expressing targets A375-NSP13 and RPMI-7951-NSP13, even at very low E:T ratio (2.5:1), but not control targets (Fig. 4 *C* and *D*), indicating that NSP13-448 specific CTLs can recognize the endogenous presented peptide of NSP13 protein. Cold target inhibition assay demonstrated that when cold targets are added, the lytic capacity of NSP13-448 specific CTLs to hot targets, A375-NSP13 and RPMI-7951-NSP13, was inhibited significantly (Fig. 4 *E* and *F*), further confirming that NSP13-448 specific CTLs lyse these targets via recognition of endogenously presented cognate peptide. ICS assays demonstrated that NSP13-448 specific CTLs produce higher levels of inflammatory cytokines, IFN-γ and TNF-α, and express higher levels of antigen-driven activation markers CD137 and CD69 when co-cultured with NSP13-448 peptide pulsed targets or NSP13 expressing targets, compared with control targets (Fig. 4*G*).

### NSP13-134 peptide specific cytotoxic T cells generated from the peripheral blood recognize SARS-CoV-2-NSP13-expressing target cells

In addition to HLA-A0101 and HLA-A0201 allele, HLA-A0301 allele covers about 22% of Caucasian and 13% of African population (27). Using the same workflow, we identified an HLA-A0301 restricted peptide, NSP13-134 (KLFAAETLK) derived NSP13 protein. Following in vitro stimulation using the ETC workflow, 11 wells of 48 wells showed clear NSP13-134 peptide tetramer positive CD8+ T cell population (Fig. S6), indicated that NSP13-134 peptide is sufficiently immunogenic to induce T cell responses in healthy donor without prior SARS-CoV-2 infection. After sorting and expansion, high purity of CD8+ and Tetramer+ NSP13-134 CTLs were generated (Fig. 5*A*). Peptide titration assay showed NSP13-134 specific CTLs were able to recognize cognate peptide as low as 100pM (Fig. 5*B*), indicating expression of relative high affinity TCR. Similar to MGP-65 and NSP13-242 specific CTLs, NSP13-134 specific CTLs were able to lyse NSP13 expressing targets Hs-578T-NSP13, even at low E:T ratio (2.5:1), but not control targets (Fig. 5*C*). Similar to MGP-65 and NSP13-242 CTL, cold target inhibition assay confirmed specific recognition of endogenously presented cognate peptide (Fig. 5*D*). ICS assay also confirmed specific recognition of NS13 expressing targets (Fig. 5*E*).

**Fig. 5.**
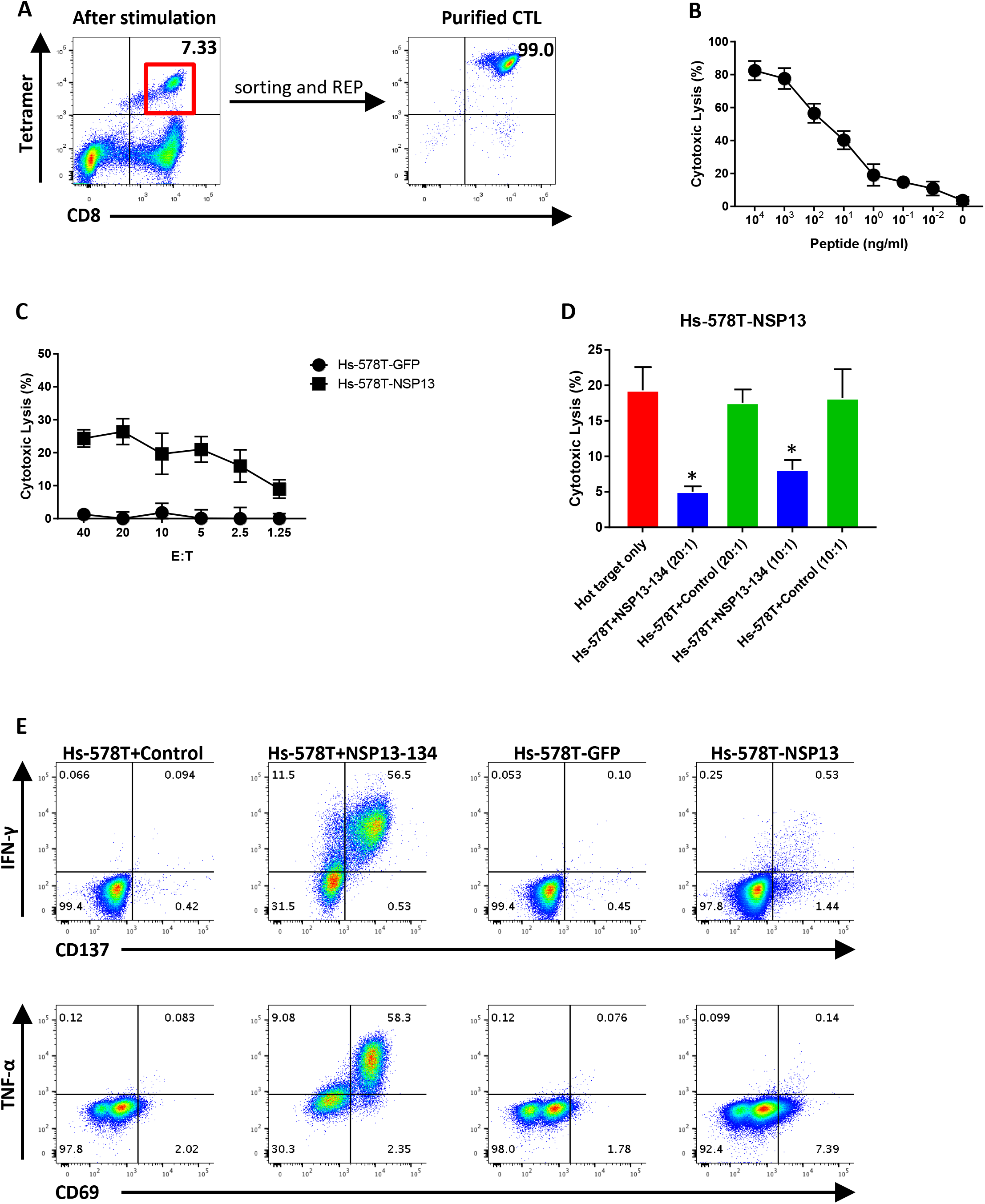
T cell generation and functional validation of non-structure protein NSP13 derived HLA-A0301 peptide NSP13-134. (A) NSP13-134 peptide (KLFAAETLK) pulsed DCs were co-cultured with autologous PBMC of HLA-A0301 healthy donor. After two rounds of stimulation, CD8+ and NSP13-134 tetramer-positive T cells were induced (left side). CD8+ and Tetramer+ T cells were sorted and then expanded with REP for two weeks to generate high purity of NSP13-134 specific CTLs (right side). (B) ^51^Cr labeled HLA-A0301 cell lines Hs-578T pulsed with various concentrations of NSP13-134 peptide were co-cultured with NSP13-134 specific CTLs at 20:1 E:T ratio. The lysis ability of NSP13-134 specific CTLs was detected with standard CRA. The data were show as average of triplicate. (C) ^51^Cr labeled NSP13 or GFP force expressing HLA-A0301 cell lines Hs-578T (Hs-578T-NSP13, Hs-578T-GFP) were co-cultured with NSP13-134 specific CTLs at various E:T ratio (from 40:1 to 1.25:1). The lysis ability of NSP13-134 specific CTLs to different targets were detected with standard CRA. The data were show as average of triplicate. (D) Cold target inhibition assay. ^51^Cr labeled Hs-578T-NSP13 cells were as hot targets. Non-radiolabeled Hs-578T cells pulsed with NSP13-134 peptide or irrelevant HLA-A0301 peptide were as cold targets. Cold target: Hot target ratio was 10:1 or 20:1. NSP13-134 specific CTLs were co-cultured with hot targets alone or hot targets together with cold targets at 20:1 E: hot T ratio. The lysis ability of NSP13-134 specific CTLs were detected with standard CRA. The data were show as average of triplicate. (E) Intracellular cytokine staining (ICS) assay. NSP13-134 specific CTLs were co-culture with Hs-578T pulsed with NSP13-134 peptide or irrelevant HLA-A0301 peptide, as well as Hs-578T-NSP13, Hs-578T-GFP at 10:1 E:T ratio in the presence of BFA for overnight. After incubation, the levels of IFN-γ and TNF-α, as well as TCR pathway down-stream activated marker CD137 and CD69 were detected using flow cytometry.

### Expansion of NSP13-400 peptide specific cytotoxic T cells from the peripheral blood of healthy donor and functional assay

In addition to HLA-A0101, HLA-A0201 and HLA-A0301, HLA-A2402 allele covers an additional of 40% Asians and 20% Caucasians (27). Using the same workflow, we identified the HLA-A2402 restricted peptide, NSP13-400 (VYIGDPAQL) of NSP13. Following in vitro stimulation using 9 of 48 wells showed clear NSP13-400 peptide tetramer positive CD8+ T cell population (Fig. S7), After sorting and expansion, high purity of CD8+ and Tetramer+ NSP13-400 CTLs were generated (Fig. 6*A*). Similar to MGP-65 CTLs, NSP13-400 specific CTLs showed very high recognition affinity for the cognate peptide in peptide titration assay, as low as 10pM concentration (Fig. 6*B*) and in accordance with peptide titration assay, very high, specific lysis of NSP13 expressing targets Hs-578T-NSP13 and M14-NSP13, even at E:T ratios as low as 1.25:1 (Fig. 6 *C* and *D*). Similarly, cold target inhibition assay showed specific level of NSP13-400 CTLs to hot target Hs-578T-NSP13 and M14-NSP13 which were significantly inhibited with addition of cold targets (Fig. 6 *E* and *F*), further confirmed that NSP13-400 specific CTLs lyse the SARS-CoV-2 targets via recognition of endogenously presented cognate peptide.

**Fig. 6.**
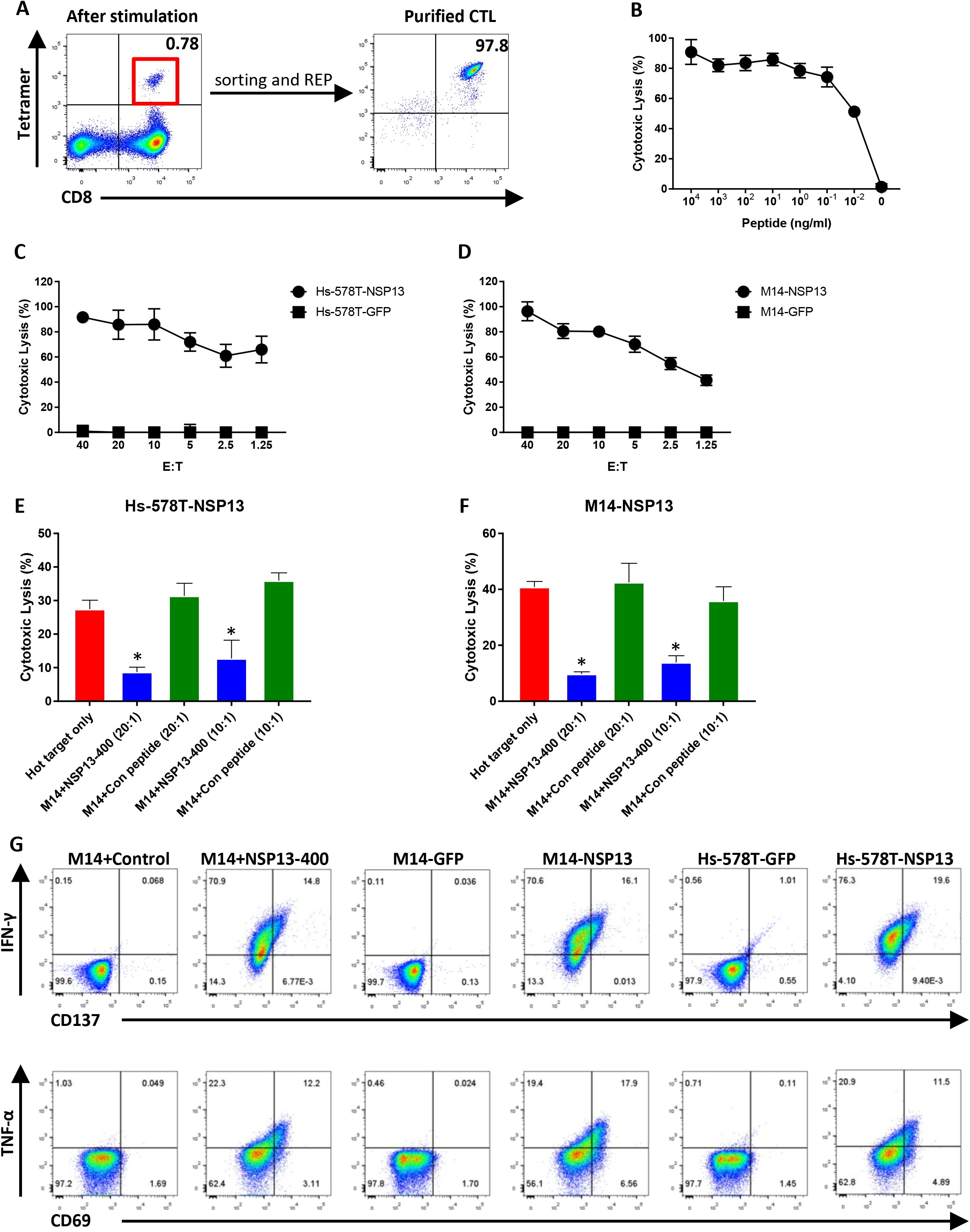
T cell generation and functional validation of non-structure protein NSP13 derived HLA-A2402 peptide NSP13-400. (A) PBMC from HLA-A2402 healthy donor were co-cultured with NSP13-400 peptide (VYIGDPAQL) pulsed autologous DCs. After two rounds of stimulation, CD8+ and NSP13-400 tetramer-positive T cells were induced (left side). After sorting and REP for CD8+ and Tetramer+ T cells with two weeks, high purity of NSP13-400 specific CTLs were expanded (right side). (B) ^51^Cr labeled HLA-A2402 cell lines M14 pulsed with various concentrations of NSP13-400 peptide were co-cultured with NSP13-400 specific CTLs at 20:1 E:T ratio. The lysis ability of NSP13-400 specific CTLs was detected with standard CRA. The data were show as average of triplicate. (C, D) ^51^Cr labeled NSP13 or GFP force expressing HLA-A2402 cell lines Hs-578T (Hs-578T-NSP13, Hs-578T-GFP) and M14 (M14-NSP13, M14-GFP) were co-cultured with NSP13-400 specific CTLs at various E:T ratio (from 40:1 to 1.25:1). The lysis ability of NSP13-400 specific CTLs to different targets were detected with standard CRA. The data were show as average of triplicate. (E, F) Cold target inhibition assay. ^51^Cr labeled Hs-578T-NSP13 and M14-NSP13 were as hot targets. Non-radiolabeled M14 cells pulsed with NSP13-400 peptide or irrelevant HLA-A2402 peptide were as cold targets. Cold target: Hot target ratio was 10:1 or 20:1. NSP13-400 specific CTLs were co-cultured with hot targets alone or hot targets together with cold targets at 20:1 E: hot T ratio. The lysis ability of NSP13-242 specific CTLs were detected with standard CRA. The data were show as average of triplicate. (G) Intracellular cytokine staining (ICS) assay. NSP13-400 specific CTLs were co-culture with M14 pulsed with NSP13-400 peptide or irrelevant HLA-A2402 peptide, as well as Hs-578T-NSP13, Hs-578T-GFP, M14-NSP13, M14-GFP at 10:1 E:T ratio in the presence of BFA for overnight. After incubation, the levels of IFN-γ and TNF-α, as well as TCR pathway down-stream activated marker CD137 and CD69 were detected using flow cytometry.

ICS assay also confirmed specific recognition of NS13 expressing targets (Fig. 6*G*). IFN-γ and TNF-α level were strikingly elevated in comparison with other SARS-CoV-2 CTL target assays suggesting high density endogenous presentation.

In summary, using our MHC IP elution and MS identification workflow, we discovered five HLA Class-I restricted peptide derived from structure protein MGP and non-structure protein NSP13 of SARS-CoV-2 presented by several HLA alleles (HLA-A0101, HLA-A0201, HLA-A0301 and HLA-A2402) which cover approximately 80% of Caucasian and Asian populations. All five peptide were highly immunogenic and capable of readily eliciting T cell responses among healthy COVID19-negative donors. All five SARS-CoV-2 specific CTLs recognize endogenously presented cognate peptide and specifically lyse SARS-CoV-2 + targets.

## Discussion

To date, nearly 1,500 predicted Class I epitopes for SARS-CoV2 have been identified by in silico prediction methods, and in some cases, ‘validated’ by eliciting T cell responses using PBMC of patients with COVID19 (4, 5). These peptides have been used extensively to evaluate the T cell response of patients, and occasionally healthy donors, to COVID19, and COVID19 vaccines, and increasingly, to develop T cell-based therapies. What has not been demonstrated however, is whether any of these 1,500 predicted peptides are in fact processed and presented by SARS-CoV2+ cells and represent naturally-occurring epitopes recognized by T cells. A preliminary screen of predicted SARS-CoV2 epitopes considered “immunodominant” among widely cited reports appears to support this premise: previously we found that 7 of these 8 predicted peptides were unable to elicit a T cell response that would lead to recognition of SARS-CoV2+ targets suggesting that responses to these peptides may be artifactual, or at best cross-reactive (4, 7, 15–22). To date, there has been no empiric validation of SARS-CoV2 epitopes for immunogenicity.

We postulate that immunogenic epitopes for SARS-CoV-2 are best defined empirically by directly analyzing peptides eluted from MHC and then validating immunogenicity by determining if such peptides can elicit T cells recognizing SARS-CoV-2 antigen-expressing targets. Mass spectrometry (MS) is an ideal analytical approach to precisely identify the naturally expressed antigenic epitopes and enables investigators to address the complexity associated with differential expression and processing of antigenic proteins by infected cells. Based on immunoaffinity capture of the MHC-antigenic peptide complex from cells engineered to express SARS-CoV2 genes, our approach allowed for direct profiling and identification of the SARS-CoV2 immunopeptidome. By eliciting T cell responses against these candidate epitopes, we confirm empiric recognition of SARS-CoV2+ cells and endogenous presentation of these peptides.

In this study, we identify and validate 5 Class I-restricted SARS-CoV2 epitopes expressed by structural (MGP) and non-structural genes (NSP13), presented by Class I alleles (HLA-A*0101, A*0201, A*0301, HLA-A*2402) prevalent among > 75% of the general population. Using recombinant vectors encoding these alleles, we engineered expression of highly conserved regions of SARS-CoV2 membrane glycoprotein (MGP) and non-structural protein-13 (NSP13) genes, recovered MHC, eluted peptides and applied data dependent analysis liquid chromatography-tandem mass spectrometry (DDA MS/MS) yielding over 12,000 spectra, which were then deconvoluted and filtered to a handful of candidate peptide epitopes. The immunogenicity of 5 peptides was validated on the basis of their ability to elicit peptide-specific T cells capable of recognizing and killing SARS-CoV2-expressing target cells.

The importance of eliciting a meaningful anti-viral T cell response has been well-documented; SARS- and MERS-responsive T cells were found to have a protective role (28). Emergence of SARS-CoV-2-specific T cell responses were recently shown to be associated with a sustained viral clearance and highlight the importance of developing vaccines that promote cellular immunity against SARS-CoV-2 (29–31).

Recently, the emergence of mutant escape variants of SARS-CoV2 has led to global concerns over possible breaches in viral protection following immunization with current vaccines which elicits a predominantly serologic response (15, 32–35). By targeting a non-surface, non-structural protein, in this case nsp13, which encodes viral helicase, escape variants are less likely to develop; in fact, none of the known variants harbor mutations among the epitope sequences identified here. Furthermore, the strategy presented allows for identification of epitopes spanning almost any SARS-CoV2 gene; selection of virus-essential gene targets provides a rational T cell-based approach to mitigate selection of antigen-loss variant and the potential for long-term viral immunoprotection.

Equally important in defining the landscape of COVID19 infection and control, its natural history, vaccine efficacy and therapeutic intervention, is an accurate measure of the SARS-CoV2-specific immune response. While the pools of predicted peptides currently in use to evaluate Class II ad Class I-restricted responses have been used extensively and appear to provide a measure of overall immune response, SARS-CoV2-specific immunity is poorly defined when the majority of peptides may not be immunogenic; the use of a highly defined subset of peptides may provide a more accurate representation of T cell immune response to COVID19 infection Although our current panel of 5 peptides is not extensive, it does represent highly conserved regions of the SARS-CoV2 genome, presented by several highly prevalent allelotypes, and may readily be applied to Class II as well as Class I-restricted epitopes. A more extensive panel of 18 epitopes from MGP, SP and NSP13 have been prepared and will be evaluated for clinical correlative studies.

## Materials and Methods

### Blood donors and Cell lines

Healthy donor peripheral blood mononuclear cells (PBMC) samples expressing the HLA-A0101, HLA-A0201, HLA-A0301 or HLA-A2402 allele were purchased from HemaCare (CA, USA) as a source of responding T cells and autologous antigen presenting cells. TAP-deficient T-B cell hybrid cell line T2, melanoma cell line A375 (HLA-A0101/0201), RPMI-7951 (HLA-A0101/0201) and package cell lines Phoenix-GP, 293T were purchased from ATCC (VA, USA). Melanoma cell lines Hs-578T (HLA-A0301/2402) and M14 (HLA-A1101/2402) were purchased from NCI. Melanoma cell line Mel624 (HLA-A0201) was the gift from Dr. Steven Rosenberg (NCI). Lymphoblastoid cell lines (LCL) are EBV-transformed lymphoblastoid cell lines established in our laboratory. Cancer cell lines were maintained in RPMI-1640 media with Hepes (25 mM), L-glutamine (4mM), penicillin (50 U/ml), streptomycin (50 mg/ml), sodium pyruvate (10 mM), nonessential amino acids (1 mM), and 10% fetal bovine serum (FBS) (Sigma, MO, USA). Phoenix-GP were cultured in DMEM media with Hepes (25 mM), L-glutamine (4mM) and 10% FBS.

### Lentivirus transduction

The cDNA of membrane glyco-protein (MGP) and Non-structure protein 13 (NSP13) of ORF1b from SARS-CoV-2 were purchased from Genscript (NJ, USA) and cloned into lentiviral vector pLVX (TAKARA, CA, USA) with fusion of GFP. In this vector, the expressing gene was driven by human EF1 promoter. MGP-pLVX or NSP13-pLVX lentiviral vector were transfected into package cell line 293T, together with package vectors contain VSVG envelop vector to make lentivirus. A375, Mel624, RPMI-7951, Hs-578T and M14 cell lines were infected with MGP-pLVX or NSP13-pLVX lentiviral vectors and the stable cell lines were screened with puromycin selection. MGP or NSP13 gene expressing efficiency was detected by analyzing the percentage of GFP using flow cytometry (NovoCyte Flow Cytometer Systems, Agilent, CA, USA).

### HLA Class-I binding peptide identification

HLA Class-I binding peptide isolation and identification via immune-precipitation (IP) and tandem mass spectrometry (MS) methods are referenced from prior study (36). Briefly, about 300 to 500 million cells engineered to express the SARS-CoV-2 MGP or NSP13 gene were homogenized in cold NP40 lysis buffer supplemented with protease inhibitor cocktail (Roche, CA, USA). Lysates were cleared by subsequent centrifugation and filtering steps. HLA class I molecules from the cleared lysate were incubated with anti-HLA-A, B, C monoclonal antibody (W6/32) coupled Sepharose-4B resin (GE Healthcare, IL, USA) at room temperature for 2 hours. The un-bound protein was washed by PBS. The HLA molecules with their bound peptides were then eluted from the affinity column with 0.1N acetic acid. The detached peptides were separated from HLA molecules using 3 kDa cut-off centrifugal ultrafilters (Millipore, MO, USA), and then concentrated using vacuum centrifugation.

Peptides were reconstituted in 0.1% formic acid in water prior to mass spectrometry acquisition. New Objective PicoFrit nanospray column (360 μm OD × 75 μm ID) was packed with Dr. Maisch 3 μm ReproSil-Pur C18 beads to 30 cm. The same C18 beads were used to pack 25mm trap column using 360 μm × OD 150 μm ID fused silica capillary fitted with Kasil on one end. Peptides were separated using Thermo Scientific *EASY-nLC* 1200. Solvent A was 0.1% formic acid in water, and solvent B was 0.1% formic acid in 80% acetonitrile. For each injection, 10 – 15 μL was loaded and eluted using 25 – 60 minute gradient from 5 to 40% solvent B at 250 – 300 nL/min. Thermo Scientific Q-Exactive HF or Orbitrap Exploris 480 tandem mass spectrometer was used to acquire mass spectra using data-dependent acquisition (DDA) or Parallel reaction monitoring (PRM).

### DDA acquisition on Q-Exactive HF

Precursor spectra (400–1600□m/z) were collected at 60,000 resolution with Automatic Gain Control (AGC) target set at 3e6 and maximum inject time of 100 ms. Fragment spectra were collected at 15,000 resolution with AGC target set at 1e5 and maximum inject time of 25□ms. The isolation width was set to 1.6□m/z. Normalized collision energy was set at 27. Top-20 most intense precursor ions for fragmentation was selected. Charge exclusion was enabled to include only precursor charges between +2 and +4 with AGC threshold of 5e3. Dynamic exclusion was set to 10 seconds to exclude all isotopes clusters.

### PRM acquisition Orbitrap Exploris 480

Precursor spectra (400–1600□m/z) were collected at 30,000 resolution with standard AGC target set and automatic maximum inject time. RF lens was set at 50%, Cycle time, at 3 seconds. Fragment spectra scan range was set at 500 – 1600 and collected at 15,000 resolution with AGC target set at standard and automatic injection time. The isolation width was set to 2 m/z with unscheduled time mode. Normalized collision energy was set at 30%. An inclusion list containing m/z values of protonated precursor peptide ions of interest was generated in Skyline-daily (version 20.2.1.135).

### Peptide selection and validation

To analyze the acquired MS/MS spectra, the spectra were searched against the Swiss-Prot protein database (version 2020_05) by using Mascot search engine node (version 2.6) within Proteome Discoverer 2.3. The searches were performed with a precursor peptide mass tolerance of 15 ppm and fragment ion mass tolerance of 15 ppm using monoisotopic parent and fragment ion masses allowing for two missed cleavages without enzyme specification. Taxonomy was restricted to Homo sapiens (20,386 sequences) and Viruses (17,008 sequences). The False Discovery Rate (FDR) was determined using “Target Decoy PSM Validator” node within Proteome Discoverer (version 2.3) and protein/peptide with an FDR of ≤ 1% being retained for further analysis. The predicted binding to these putative peptides was determined and filtered by IEDB T Cell Epitope Prediction Tools (http://tools.iedb.org/main/tcell/).

### Generation of SARS-CoV-2 specific T cells

Generation of antigen-specific T cell stimulation was performed according to our endogenous T cell (ETC) generation workflow (37). Briefly, adherent PBMCs were treated with GM-CSF (800 U/mL) and IL-4 (500 U/mL) for 6 days to generate immature DC (iDC), and the iDC were then matured with a cytokine cocktail containing TNF-α (10 ng/mL), IL-1β (2 ng/mL), IL-6 (1000 U/mL), PGE-2 (1000 ng/mL) for an additional 2 days. Candidate SARS-CoV-2 peptide was pulsed on HLA-matched mature DC for MGP-65 (FVLAAVYRI, HLA-A0201), NSP13-448 (IVDTVSALVY, HLA-A0101), NSP13-242 (TLVPQEHYV, HLA-A0201), NSP13-134 (KLFAAETLK, HLA-A0301), NSP13-400 (VYIGDPAQL, HLA-A2402) (all purchased from Genscript, NJ, USA) in PBS/HSA. The peptide pulsed DCs were then co-cultured with autologous PBMC in RPMI-1640 with Hepes (25 mM), L-glutamine (4mM), penicillin (50 U/ml), streptomycin (50 mg/ml), sodium pyruvate (10 mM), and 10% human AB serum. After 7 days in culture, T cell cultures were restimulated with peptide-pulsed DC as before. IL-2 (10 U/mL) and IL-7 (5 ng/mL) were added on the second day.

### Sorting and expansion

After two stimulation cycles, an aliquot of each well was stained with custom PE-conjugated MHC tetramer folded with HLA matched SARS-CoV2 peptide, and with APC-Cy7 conjugated anti-CD8 antibody (Biolegend, CA, USA). Cells were washed and analyzed by flow cytometry (NovoCyte Flow Cytometer Systems, Agilent, CA, USA). The tetramer positive staining wells were pooled and CD8/Tetramer double positive population were sorted using flow cytometric sorting (ARIA II sorter, BD, CA, USA) and then expanded using a rapid expansion protocol (REP) in a sterile 25 mL flask containing RPMI-1640 with Hepes (25 mM), L-glutamine (4mM), penicillin (50 U/ml), streptomycin (50 mg/ml), sodium pyruvate (10 mM), 10% fetal bovine serum (FBS), irradiated PBMC and LCL feeder cells, as previously described (38). After expansion, the purity of antigen specific T cells were determined with anti-CD8 antibody and MGP-65, NSP13-448, NSP13-242, NSP13-134, or NSP13-400 tetramer staining again.

### Function analysis of SARS-CoV-2 specific T cells

The cytotoxicity of purified SARS-CoV-2 specific T cells following expansion was confirmed using standard chromium (^51^Cr) release assay (CRA). Peptide dose titration experiments were performed to test cognate peptide recognition of SARS-CoV-2 CTL. Titrating concentrations of MGP-65 or NSP13-242 peptide-pulsed T2 cells (HLA-A2+) were used for evaluating MGP-65 or NSP13-242 CTL; NSP13-448 peptide pulsed A375 cells (HLA-A1+), for NSP13-448 CTL; NSP13-134 peptide pulsed Hs-578T cells (HLA-A3+) for NSP13-134 CTL, and NSP13-400 pulsed M14 cells (HLA-A24+) for NSP13-400 CTL. Target cells were labeled with 100 μCi ^51^Cr (Perkin Elmer, CA, USA) in 1 ml of tumor cell culture media for 1 hour, then washed and plated at 2, 000 target cells per well in triplicate. MGP-65, NSP13-448, NSP13-242, NSP13-134 or NSP13-400 specific T cells were added at effector-to-target (E:T) of 20:1 cell ratio for 4 hours. Supernatant was collected from the wells and ^51^Cr measured with a gamma radiation counter. The percentage of specific target cell lysis was calculated, correcting for background ^51^Cr release and relative to a maximum ^51^Cr release as measured by NP40 lysed target cells (39).

Tumor cell lines engineered to express MGP or NSP13 genes were used as targets to evaluate SARS-CoV-2 specific T cell recognition of endogenously presented epitopes. A375-MGP, Mel624-MGP (HLA-A0201+, MGP+), A375-GFP, Mel624-GFP (HLA-A0201+, GFP+), were used to evaluate MGP-65 specific T cell activity; A375-NSP13, RPMI-7951-NSP13 (HLA-A0101+, MGP+), A375-GFP, RPMI-7951-GFP (HLA-A0101+, GFP+), for NSP13-448 specific T cell activity; A375-NSP13, Mel624-NSP13 (HLA-A0201+, NSP13+), A375-GFP, Mel624-GFP, for NSP13-242 specific T cell activity; Hs-578T-NSP13 (HLA-A0301+, HLA-A2402+, NSP13+), Hs-578T-GFP (HLA-A0301+, HLA-A2402+, GFP+), for NSP13-134 specific T cell activity and M14-NSP13 (HLA-A2402+, NSP13+), Hs-578T-NSP13, M14-GFP (HLA-A2402+, GFP+), Hs-578T-GFP, for NSP13-400 specific T cell activity. ^51^Cr labeled target cells were co-cultured with SARS-CoV-2 specific T cells at varying effector-to-target (E:T) cell ratios. After the incubation period, antigen-specific target cell lysis was determined as above.

### Cold target inhibition assay

To confirm, epitope and antigen-specificity against relevant tumor targets, cold target inhibition assays were performed as described previously (40). For MGP-65 specific T cell test, A375-MGP and Mel624-MGP cells labeled with ^51^Cr were used as ‘hot’ targets. Non-radiolabeled T2 cells pulsed with MGP-65 peptide (10μg/ml) were used as cold targets. Non-radiolabeled T2 cells pulsed with M26 control peptide (ELAGIGILTV, HLA-A0201) were used as control cold targets. Before co-culturing the MGP-65 specific T cells with hot target, cold targets (at 10- or 20-fold greater numbers than radiolabeled hot targets) were added and incubated with a given antigen-specific T cells for one hour. ^51^Cr-labeled hot targets were added at E:T ratio (20:1) and incubated for further 4 hours. After the incubation period, target cell lysis by MGP-65 specific T cells was determined above. Similarly, for NSP13-448 specific T cells, A375-NSP13 and RPMI-7951-NSP13 cells labeled with ^51^Cr were used as hot targets. Non-radiolabeled A375 cells pulsed with NSP13-448 peptide or VGLL1 HLA-A0101 peptide (LSELETPGKY) (36) were used as cold targets and control cold targets, respectively. For NSP13-242 specific T cells, A375-NSP13 and Mel624-NSP13 cells labeled with ^51^Cr were used as hot targets. Non-radiolabeled T2 cells pulsed with NSP13-242 peptide or M26 peptide were used as cold targets and control cold targets, respectively. For NSP13-134 specific T cells, Hs-578T-NSP13 cells labeled with ^51^Cr were used as hot targets. Non-radiolabeled Hs-578T cells pulsed with NSP13-134 peptide or A3 control peptide (KVFPCALINK, HLA-A0301) were used as cold targets or control cold targets. For NSP13-400 specific T cell test, Hs-578T-NSP13 and M14-NSP13 cells labeled with ^51^Cr were used as hot targets. Non-radiolabeled M14 cells pulsed with NSP13-400 peptide or MAGEA4 HLA-A2402 peptide (NYKRCFPVI) were used as cold targets or control cold targets.

### Intracellular staining (ICS) assay

One million SARS-CoV-2 specific T cells (MGP-65, NSP13-448, NSP13-242, NSP13-134, NSP13-400) were co-cultured with 1×10^5^ relevant target cells (10:1 E:T ratio) overnight in the presence of Brefeldin A (BFA) (Biolegend, CA, USA), and the next day, stained with APC-Cy7 conjugated anti-CD8 antibody (Biolegend, CA, USA). After washing, the cells were fixed and permeabilized with Intracellular Fixation & Permeabilization Buffer Set (eBioscienceTM, NY, USA), and then stained with APC conjugated anti-CD137, FITC conjugated anti-CD69, PE conjugated anti-IFN-γ, Pacific blue conjugated anti-TNF-α antibody (all purchased from Biolegend, CA, USA). After washing, the expressing level of CD137, CD69, IFN-γ and TNF-α were determined using flow cytometry assay (LSRFortessa X-20 Analyzer, BD, CA, USA).

### Statistical analysis

Data analysis was performed using GraphPad Prism version 7.03. Normally distributed data were analyzed using parametric tests (ANOVA or unpaired t test). All *p* values are indicated in figure legends. Flow Cytometry data were analyzed using FlowJo (version 10).

## Acknowledgments

We would like to thank the help from South Campus Flow Cytometry & Cell Sorting Core of UT MD Anderson Cancer Center, which is supported by NCI P30CA016672. This work was supported by National Institutes of Health grant RO1 3R01CA237672-02S1 (to C.Y.) and the Parker Institute of Cancer Immunotherapy (C.Y.)

## Author Contributions

K.P., Y.C., C.Y. conducted overall experimental design. K.P. established SARS-CoV-2 MGP and NSP13 overexpressing cell lines and performed the MHC/peptide elution. E.H., M.M. conducted mass spectrometry detection for MHC/peptide. Y.C. performed the analysis of the MS data, screening of the candidate peptide and sequence blast of the peptide. K.P., M.C., J.W., I.L., R.S., S.S. conducted SARS-CoV-2 MGP and NSP13 specific CTL generation and functional validation. K.P., Y.C., S.S., C.Y. wrote the paper. C.Y. was responsible for the supervision of laboratory studies and manuscript writing and editing.

