## Supplemental Figures for "Immunogenic SARS-CoV2 Epitopes Defined by Mass Spectrometry"

### Slide 1
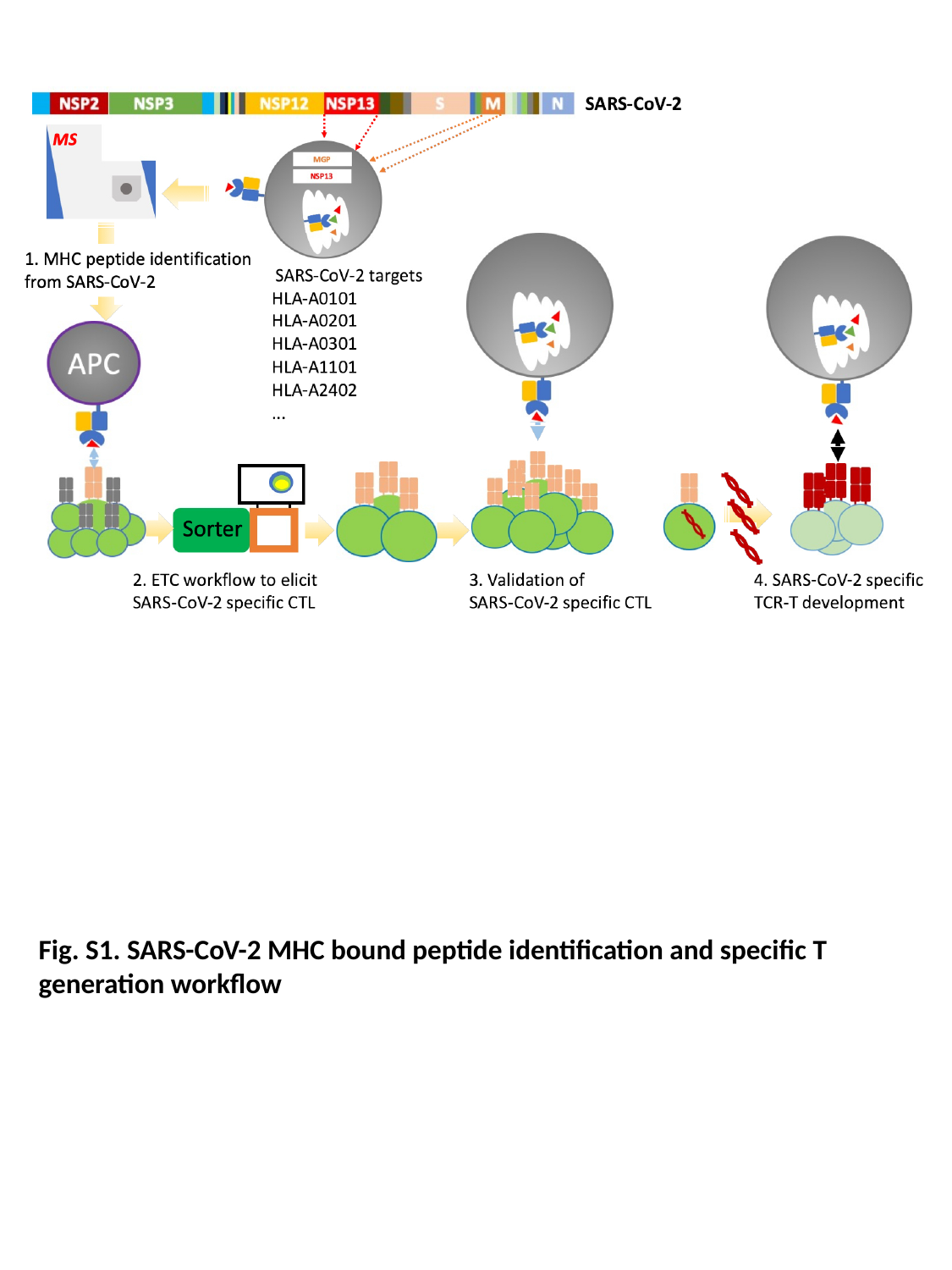

Fig. S1. SARS-CoV-2 MHC bound peptide identification and specific T generation workflow

### Slide 2
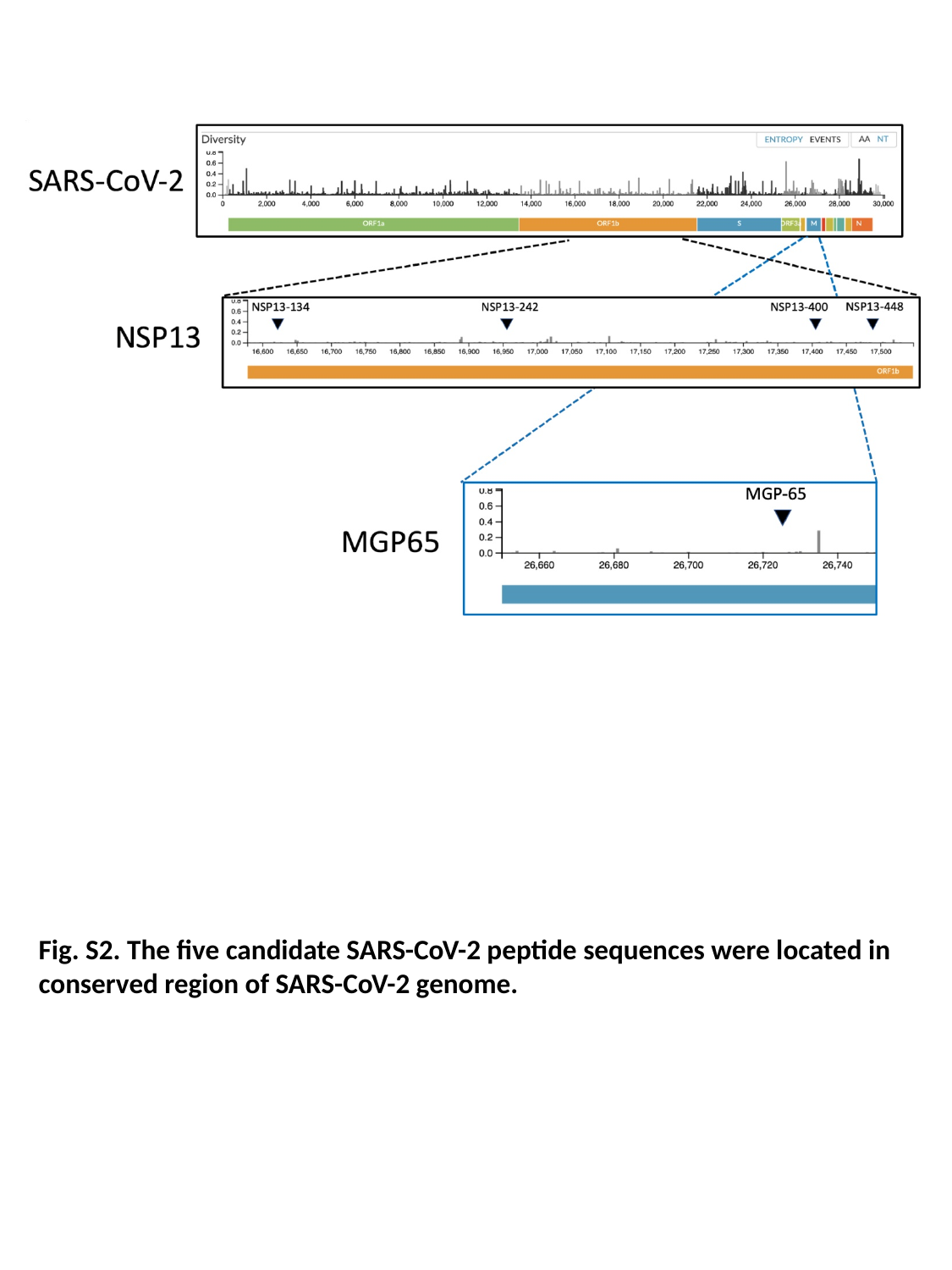

Fig. S2. The five candidate SARS-CoV-2 peptide sequences were located in conserved region of SARS-CoV-2 genome.

### Slide 3
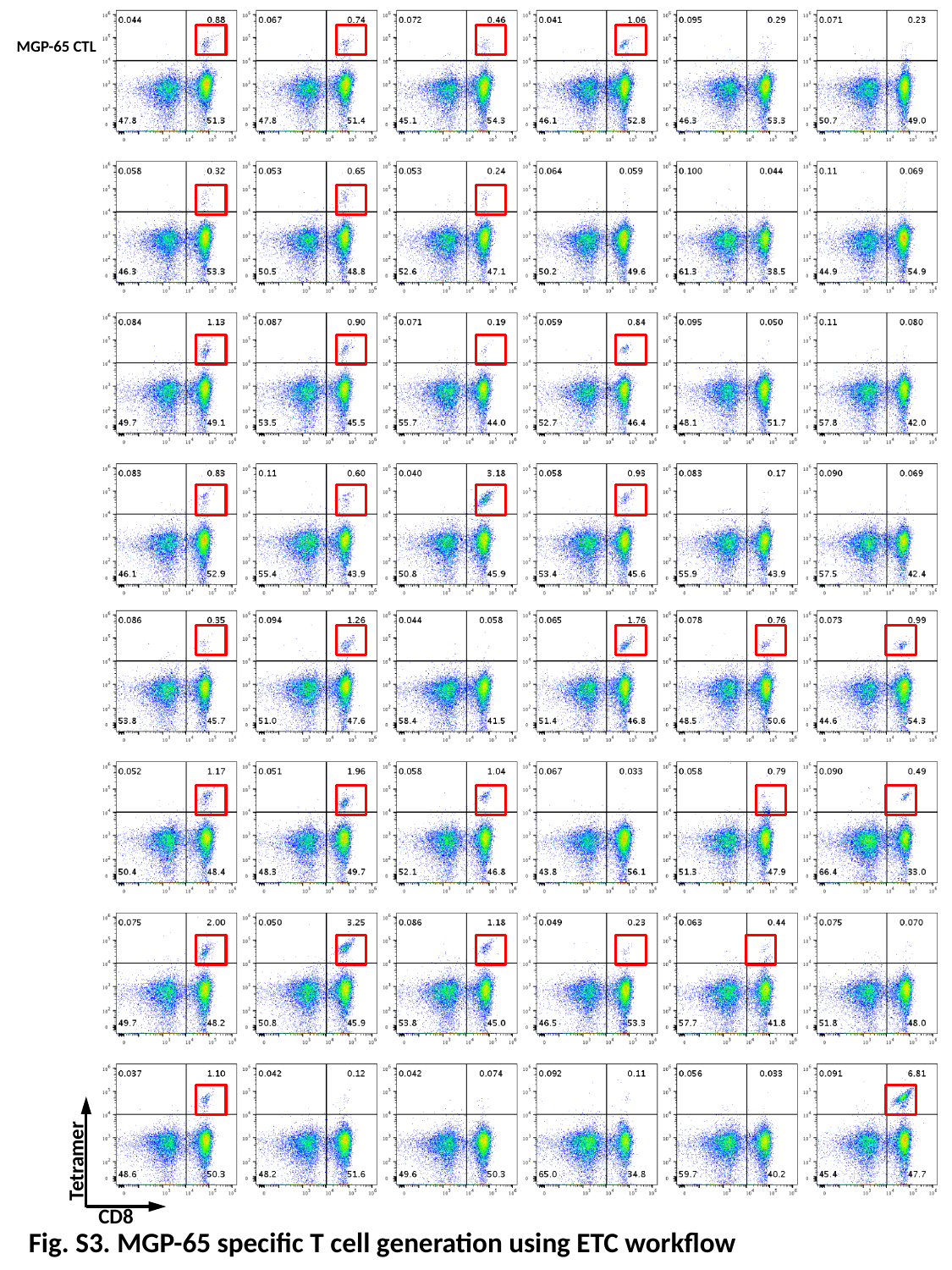

MGP-65 CTL
Tetramer
CD8
Fig. S3. MGP-65 specific T cell generation using ETC workflow

### Slide 4
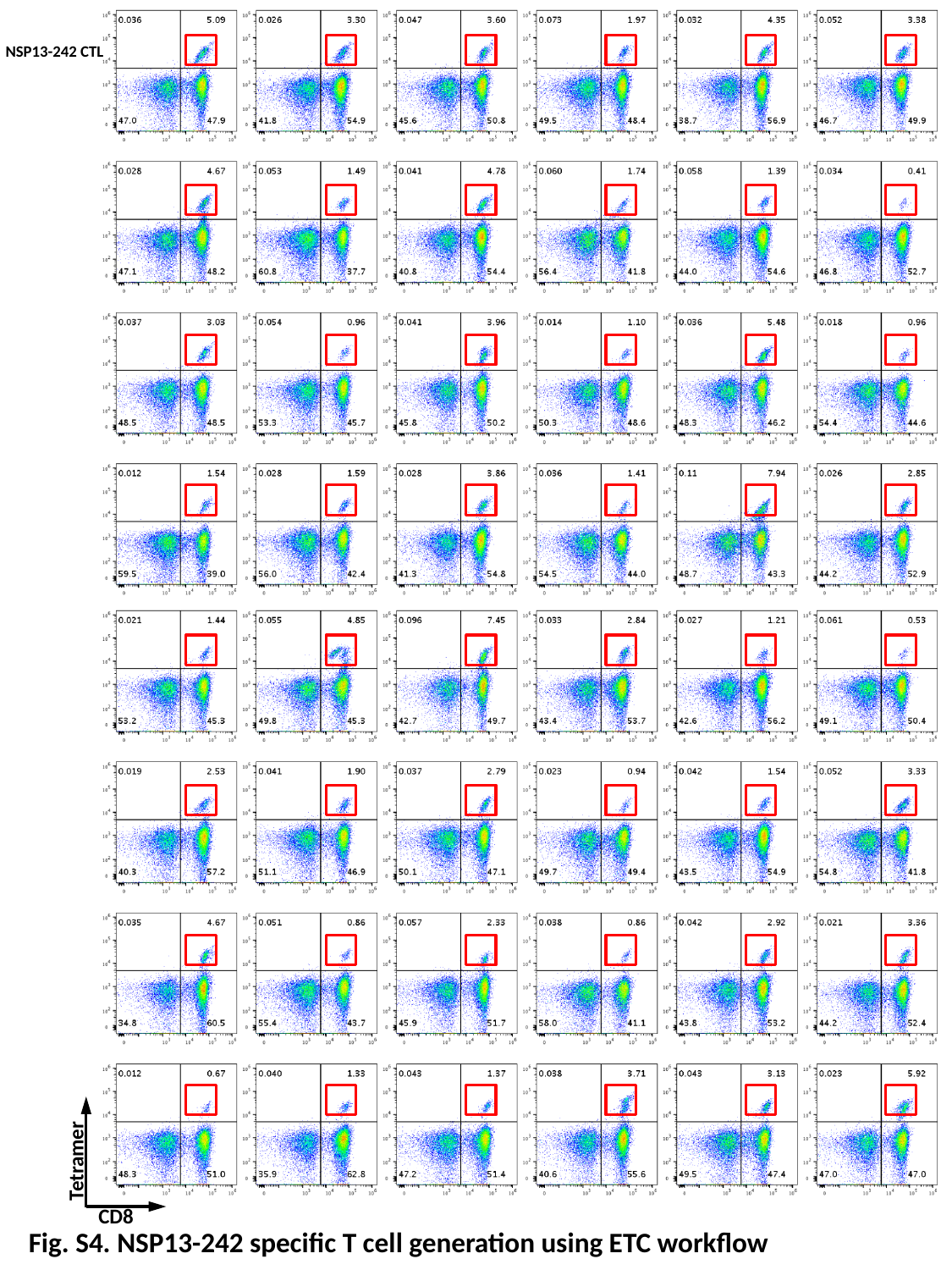

NSP13-242 CTL
Tetramer
CD8
Fig. S4. NSP13-242 specific T cell generation using ETC workflow

### Slide 5
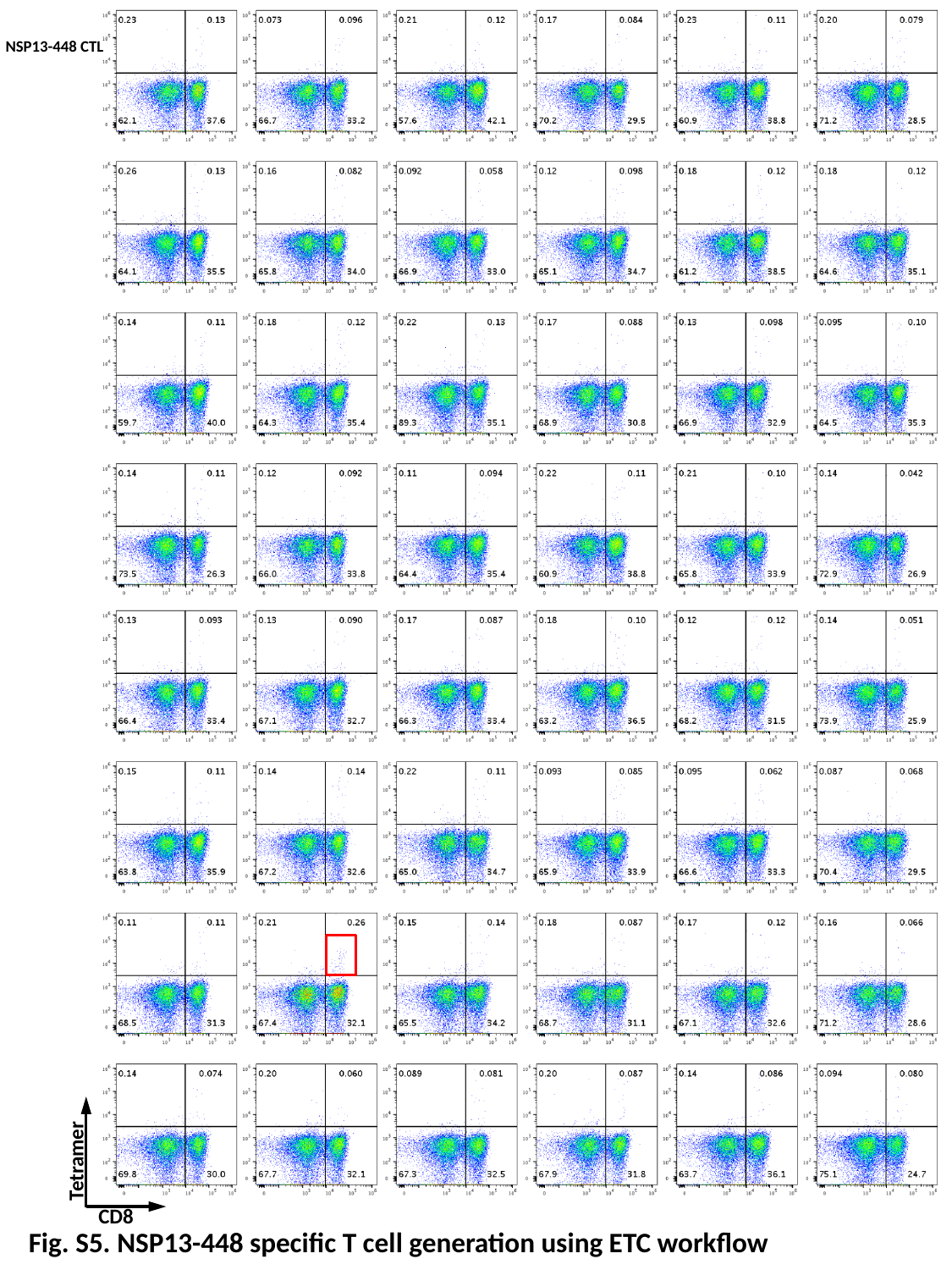

NSP13-448 CTL
Tetramer
CD8
Fig. S5. NSP13-448 specific T cell generation using ETC workflow

### Slide 6
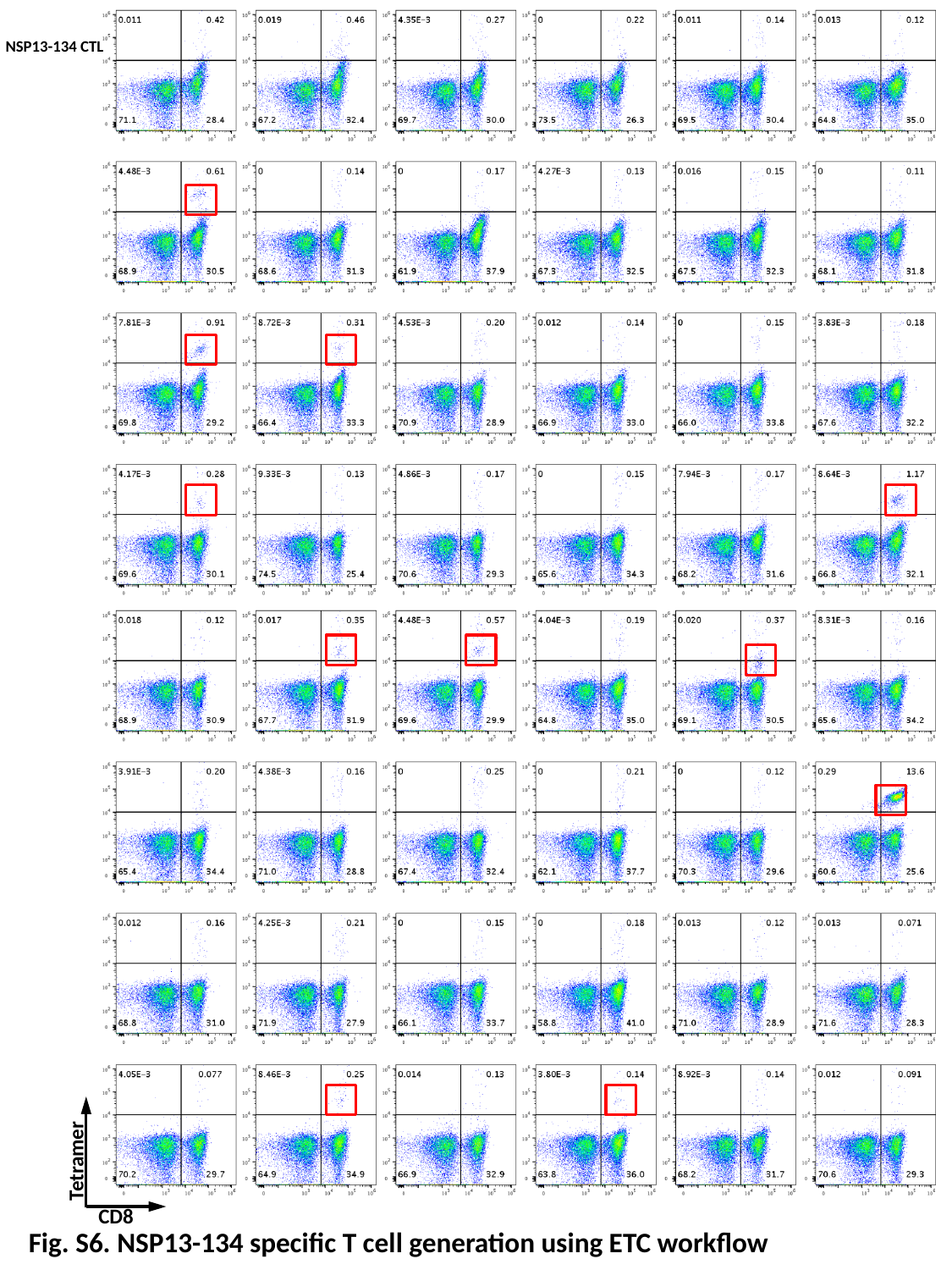

NSP13-134 CTL
Tetramer
CD8
Fig. S6. NSP13-134 specific T cell generation using ETC workflow

### Slide 7
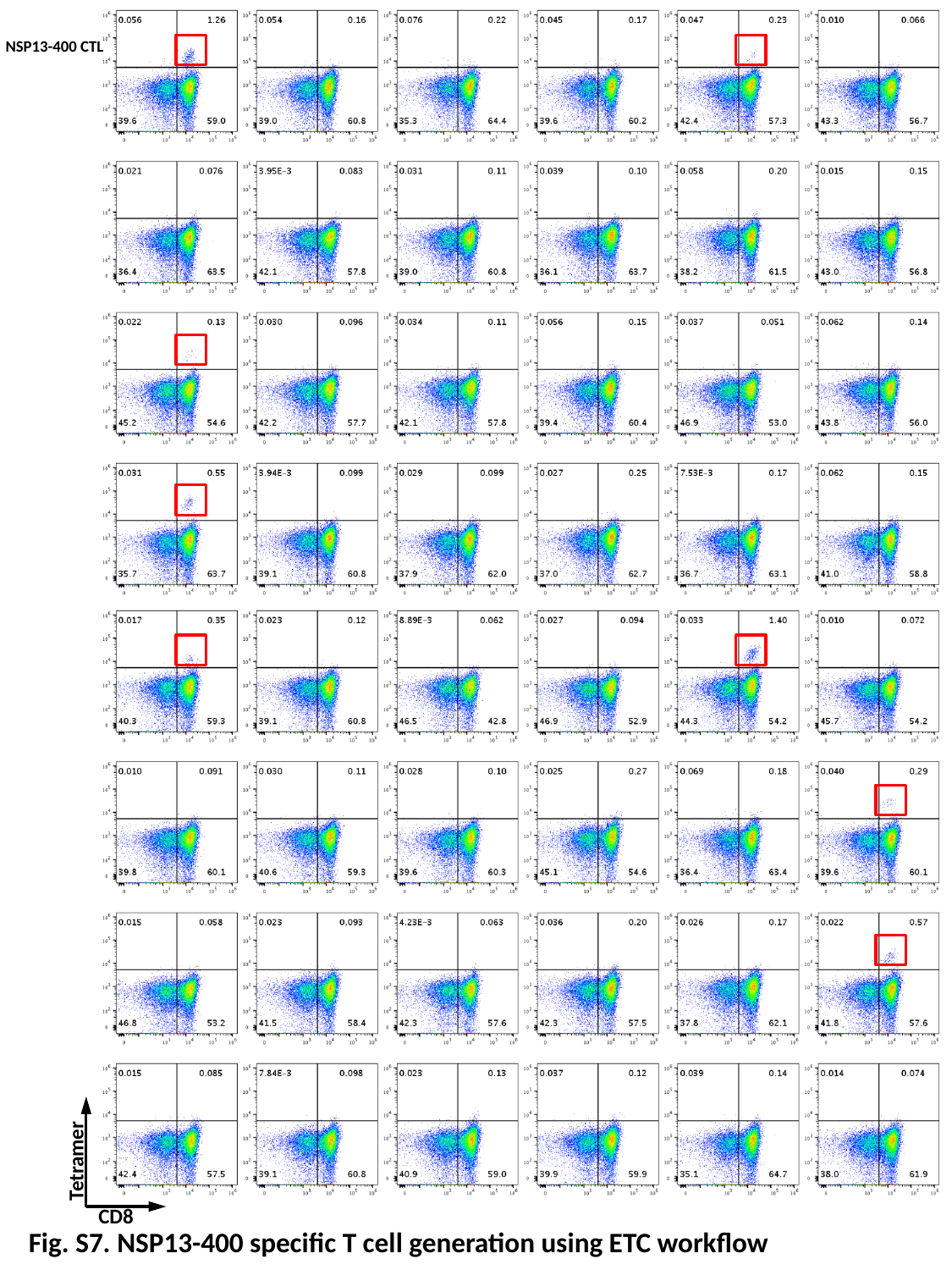

NSP13-400 CTL
Tetramer
CD8
Fig. S7. NSP13-400 specific T cell generation using ETC workflow
