## Supplemental Table-1 for "Immunogenic SARS-CoV2 Epitopes Defined by Mass Spectrometry"

**Supp-Table-1. Predicted high HLA binding peptide of MGP or NSP13 for PRM-MS analysis**

| **MGP predicted HLA-A0201 peptide** | | **NSP13 predicted HLA-A0101 peptide** | | **NSP13 predicted HLA-A0201 peptide** | | **NSP13 predicted HLA-A0301 peptide** | |
| --- | --- | --- | --- | --- | --- | --- | --- |
| **Peptide** | **Binding affinity (nM)** | **Peptide** | **Binding affinity (nM)** | **Peptide** | **Binding affinity (nM)** | **Peptide** | **Binding affinity (nM)** |
| GLMWLSYFI | 4.5 | VTDVTQLYL | 128.9 | KLVLSVNPYV | 12.3 | VVYRGTTTYK | 8.2 |
| FLYIIKLIFL | 5 | TVDSSQGSEY | 167.7 | KLSYGIATV | 20.3 | KLFAAETLK | 9.9 |
| LIFLWLLWPV | 5.6 | FAIGLALYY | 285.4 | KLNVGDYFV | 22.1 | MSYYCKSHK | 24.6 |
| VATSRTLSY | 9.8 | DSSQGSEYDY | 301.7 | SMATNYDLSV | 55.6 | QLYLGGMSYY | 37 |
| TLACFVLAAV | 12.9 | ATEETFKLSY | 348 | TLEQYVFCTV | 61.5 | SVVNARLRAK | 48.3 |
| KLLEQWNLV | 15.8 | IVDTVSALVY | 356.9 | TLVPQEHYV | 72.8 | SAQCFKMFYK | 73 |
| FLWLLWPVTL | 20.9 | KSAQCFKMFY | 555.3 | TLVPQEHYVRI | 78 | YLGGMSYYCK | 81.9 |
| LLWPVTLACFV | 34.8 | HFAIGLALYY | 577.5 | YVFCTVNAL | 93 | RFNVAITRAK | 83.8 |
| FVLAAVYRI | 38.1 | TCDWTNAGDY | 588.2 | GLALYYPSA | 104.1 | YVFTGYRVTK | 98.6 |
| LMWLSYFIA | 43.1 | CTERLKLFA | 630.1 | FKVNSTLEQYV | 108.7 | CIRRPFLCCK | 102.8 |
